# Warming Reduces Cold Hardiness of Boreal Plants but Damage Risk Varies by Species and Season

**DOI:** 10.64898/2026.05.15.725179

**Authors:** Francisco Campos-Arguedas, Erica Kirchhof, Michael North, Kyle J. Pearson, Mark P. Guilliams, Paul J. Hanson, Al P. Kovaleski

## Abstract

Winter warming is altering plant exposure to cold events, and its effects on seasonal cold hardiness dynamics remain poorly understood. Here we quantified the effect of whole-ecosystem warming (+0.00°C to +9.00°C) on bud cold hardiness of two overstory (*Larix laricina* and *Picea mariana*) and two understory (*Chamaedaphne calyculata* and *Rhododendron groenlandicum*) boreal peatland forest species across four dormant seasons. Warming reduced cold hardiness in fall and spring by delaying acclimation and advancing deacclimation, yet damage risk increased only in late winter and spring for three species. Warming also reduced snow cover, increasing temperature variability and cold damage for understory shrubs. Together, these results show that warming restructures rather than uniformly increasing cold hardiness risk, concentrating vulnerability at transitional seasons through species-specific dynamics and microclimate exposure.

## Main

Low temperature is a primary constraint on plant survival and distribution, particularly in temperate and boreal regions where winter conditions exert strong selective pressure on plant communities (*1*–*8*). To persist in these environments, woody plants prepare for winter conditions by progressively increasing the cold hardiness of overwintering tissues through a dynamic physiological response that integrates daylength, and short- and long-term temperature cues. Midwinter cold hardiness and deacclimation dynamics present species- and genotype-specific limits that are tightly linked to the climate of origin as strongly temperature-dependent phenotypes (*9*–*13*). Climate change disrupts these relationships through rising temperatures and increasingly irregular seasonal temperature cycles (*14*–*16*). Experimental evidence of how climate warming will affect plants during winter remains limited, particularly at a whole-ecosystem level, and the extent to which phenotypic plasticity of species can buffer the effects of climatic changes on winter survival remains poorly understood.

Cold hardiness – the lowest temperature a tissue or plant can survive (i.e. greater cold hardiness means more negative values of temperature) – is not a static trait. The cold hardiness at any moment during the dormant season reflects continuous processes of acclimation and deacclimation that depend on genotypic and environmental conditions such as short-term and long-term thermal exposure, both of which also affect dormancy status (*17*–*19*). As a result, a plant’s cold hardiness follows a broadly predictable annual cycle of acclimation, maintenance, and deacclimation that is nonetheless sensitive to weather variability, such that vulnerability to cold damage shifts continuously through the dormant season (*14, 17, 19, 20*). This dynamic nature means that the same absolute temperature can be harmless at one point during the dormant season and lethal at another, depending entirely on the physiological state of the plant at the time of exposure.

Cold injury is often related to temperature fluctuations rather than the absolute winter minimum. Warm winter conditions can cause early deacclimation or budbreak, resulting in damage from returning deep cold or late freezes, respectively. Both instances reduce performance and survival even when long-term thermal conditions remain within the tolerance range for a species (*14, 21*–*24*). Furthermore, species with carbon-acquisitive strategies tend to break bud earlier to capture light and nutrients ahead of competitors, but do so at greater risk of late freeze damage, whereas species with more conservative strategies break bud later and more reliably avoid damaging freeze events, albeit at the cost of a shorter growing season (*25, 26*). The greater risk can be costly as cold injury also affects carbon acquisition by delaying canopy establishment following damage (*14, 27*), with effects that can span multiple seasons (*21*). Snow cover affects cold hardiness dynamics as a thermal buffer, insulating both aboveground bud tissue and shallow roots from extreme temperature fluctuations, such that plant tissues beneath snow experience a narrower range of temperatures than those above snow (*28*). While this buffering effect is relevant across perennial species, understory plants are particularly vulnerable when snow is absent or lost early due to warming, as the entire plants (roots and canopy) become exposed to a wider range of temperature fluctuations (*28, 29*). Adaptive cold hardiness regulation is therefore critical not only for preventing episodic tissue damage, which is governed by within-year temperature regimes and microclimate conditions, but also for determining long-term survival and adaptation of woody perennial plants, and their contribution to the carbon cycle.

Here, we assessed seasonal cold hardiness responses over four winter seasons (2021-2025) at the SPRUCE (Spruce and Peatland Responses Under Changing Environments) experiment in northern Minnesota (https://mnspruce.ornl.gov). At SPRUCE, a boreal peatland community is exposed to continuous warming across five target levels (+0.00°C, +2.25°C, +4.50°C, +6.75°C, and +9.00°C above ambient temperature) using open-top chambers, at either atmospheric or elevated CO_2_ (+500 ppm) (*30*). We examined temporal changes in bud cold hardiness as a response to the different environmental conditions for four occurring species from two functional groups: a deciduous conifer (larch, *Larix laricina* (Du Roi) K. Koch) and an evergreen conifer (black spruce, *Picea mariana* (Mill.) B.S.P.) make up the overstory, and two evergreen shrubs (leatherleaf, *Chamaedaphne calyculata* (L.) Moench; and Labrador tea, *Rhododendron groenlandicum* (Oeder) Kron & Judd) represent understory species. We quantified how warming and CO_2_ enrichment differentially alter seasonal cold hardiness at two levels: across contrasting functional groups (overstory and understory), and between species within the same functional group but different foliage habits (deciduous and evergreen). Together, these comparisons allow us to assess how whole-ecosystem warming reshapes the thermal limits of winter survival across a boreal plant community.

### Warming results in uneven effects depending on plant functional group

Target warming treatments at the SPRUCE experiment are calibrated based on 2m air temperature measurements taken on a non-manipulated enclosure (+0.00°C) (*30*), resulting in evenly distributed temperature differentials across treatments at tree canopy height (**Fig. 1a,b, S1a**). However, conditions were considerably more variable during dormant seasons at the level of understory species due to changes in snow cover dynamics across the season (**Fig. 1c,d, S1b**). Snow cover acted as a long-lasting insulating layer that buffered surface temperatures in colder treatments when compared to 2m temperature. However, in more highly warmed plots, lack of snow cover for understory plants resulted in exposure to a wider range of temperatures, with more negative temperatures during cold events and higher temperatures during warm events, compared to snow-covered plants in less warmed plots (**Fig. S2, S3**). Due to the snow cover dynamics, thermal regimes differed considerably between overstory and understory species. Therefore, all analysis used temperatures recorded at the relevant height for each functional group (2m for overstory and surface level for understory), rather than the target warming treatments assigned to each enclosure.

**Figure 1.**
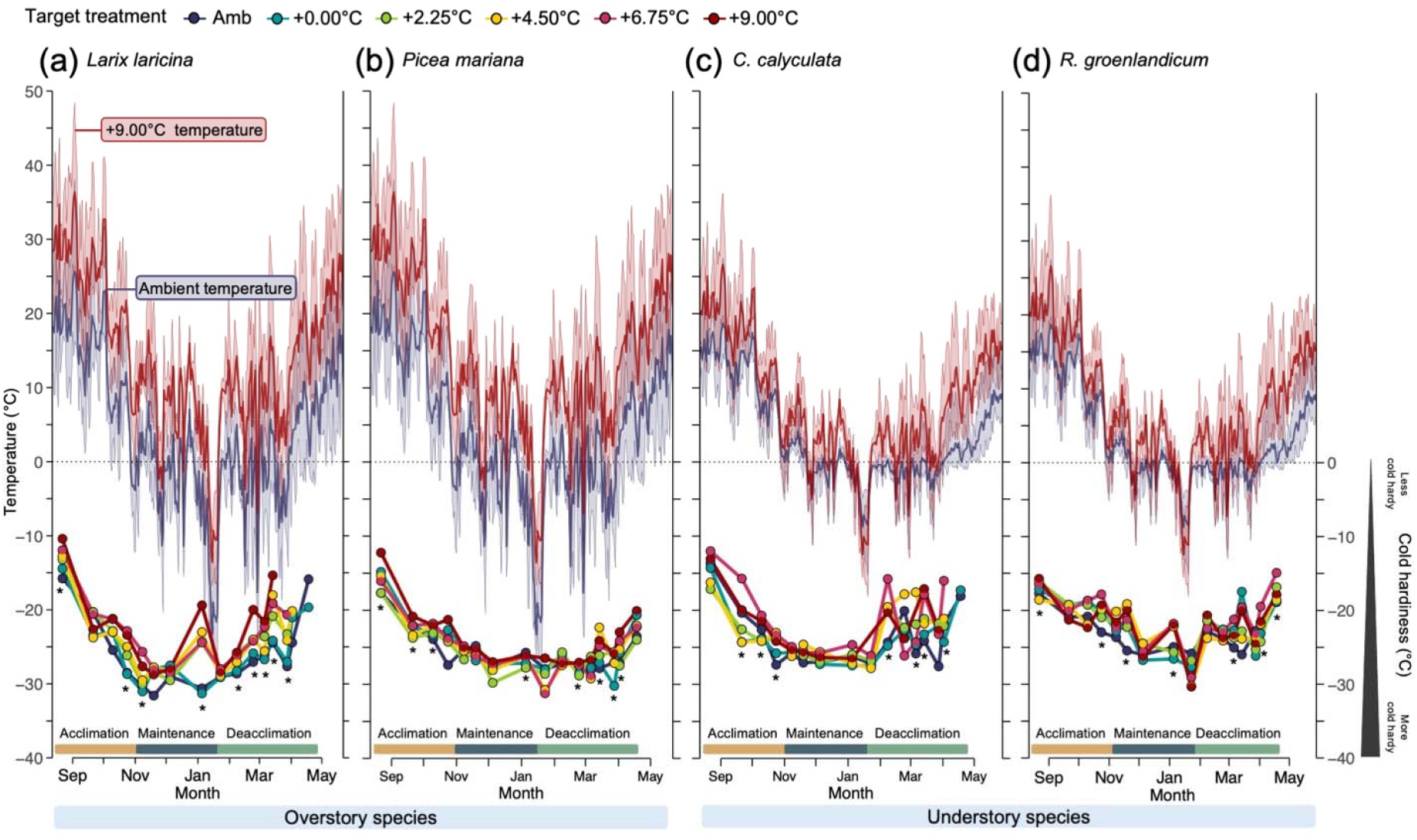
Seasonal patterns of bud cold hardiness under different warming regimes. Mean bud cold hardiness trajectories and temperature conditions during the 2023–2024 season for *Larix laricina* (**a**), *Picea mariana* (**b**), *Chamaedaphne calyculata* (**c**), and *Rhododendron groenlandicum* (**d**). Mean daily temperatures (solid lines) and minimum and maximum temperatures (ribbons) are shown for the most extreme conditions: ambient and +9.00°C target warming. Temperature data for *Larix laricina* and *Picea mariana* reflect 2m temperature measurements, while temperatures for *Chamaedaphne calyculata* and *Rhododendron groenlandicum* reflect surface-level measurements capturing the near-ground microclimate relevant to understory shrubs. Asterisks (*) indicate collections where cold hardiness is significantly affected by warming based on ΔT (*p* < 0.05). Seasonal trajectories for all four seasons are shown in **Fig. S3.**

Seasonal cold hardiness patterns were consistent across years in all four species (**Fig. 1, S3**). Overstory and understory species gained cold hardiness with decreasing temperatures in the fall, but with apparent delays due to the warming treatments. In midwinter, there are generally no significant differences across treatments, with exception of periods of considerable warm weather. Warming advanced loss of cold hardiness in the spring, particularly in overstory species. More erratic cold hardiness responses were observed in understory species, where warming caused snow cover loss that resulted in warmer treatments experiencing lower temperatures than ambient (e.g., temperatures in January 2024 – **Fig. 1c,d**). Elevated CO_2_ had no significant effect on cold hardiness (**Notes S1**), and therefore we explore here only the effect of temperature. Overall, cold hardiness presents three general phases in which we explored inter-collection and cumulative intra-seasonal thermal responsiveness: acclimation (∼mid-August to 31 October), maintenance (1 November to 28/29 February), and deacclimation (1 March to budbreak).

### Short-term thermal responsiveness is phase dependent

To evaluate short-term thermal responsiveness across physiological phases, we quantified the relationship between changes in bud cold hardiness between consecutive sample collection dates (ΔCH) and the inter-collection mean air temperature (**Fig. 2**). We characterized the distribution of observations within each phase using phase-specific centroids and bivariate 95% confidence ellipses. Each phase occupied a distinct region of the thermal and cold hardiness space (i.e., different boundaries of ellipses in the horizontal and vertical axis, respectively), with broadly consistent patterns in transition through phases across overstory and understory species.

**Figure 2.**
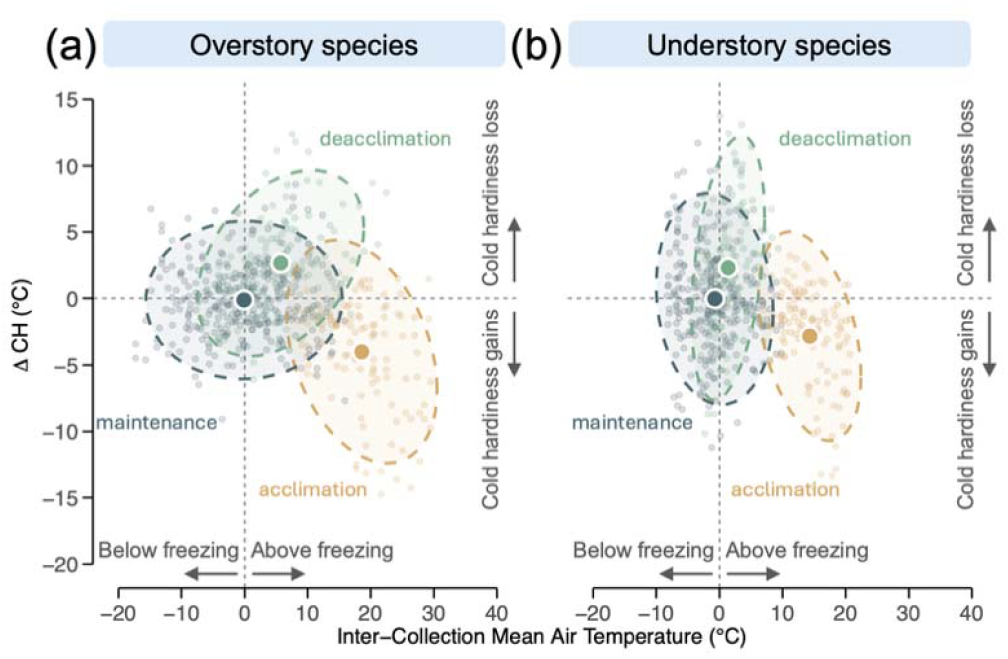
Short-term thermal responsiveness of cold hardiness across physiological phases. Relationships between inter-collection mean air temperature and short-term changes in bud cold hardiness (ΔCH) for overstory (**a**) and understory species (**b**). Negative ΔCH values indicate hardiness gain (acclimation) and positive values indicate hardiness loss (deacclimation) between consecutive sample collection points. Inter-collection mean air temperature is the average temperature between consecutive sample collection points. Points represent individual observations during acclimation (golden), midwinter maintenance (dark teal), and deacclimation (sage green) phases; filled circles indicate phase centroids and dashed ellipses represent 95% confidence regions for each phase. Treatment-specific responses are shown in **Fig. S4**.

During acclimation, buds consistently gained cold hardiness (negative ΔCH) despite experiencing warmer inter-collection temperatures than during other phases (**Fig. 2**). This reflects the late summer and early fall thermal environment under which most cold hardiness acquisition occurs (**Fig. 1, S3**). During midwinter maintenance, ΔCH values clustered near zero across a wide temperature range. This indicates that once buds reached their approximate maximum hardiness, short-term temperature fluctuations have little influence on hardiness status – a pattern consistent across both functional groups. During deacclimation, buds lost cold hardiness with rising spring temperatures. Warmer treatments amplified this loss more than they affected fall acclimation or midwinter maintenance (**Fig. S4**), demonstrating that spring warming has a disproportionate effect on cold hardiness dynamics over an annual cycle. Notably, cold hardiness loss during deacclimation occurred at lower temperatures than those driving hardiness gain during acclimation, reflecting a fundamental asymmetry in the thermal control of seasonal hardiness dynamics (**Fig. 2**). In particular, understory species showed a greater range of ΔCH within a narrower range of above-freezing inter-collection temperatures during deacclimation, showing the higher responsiveness of cold hardiness trajectories to temperature compared to overstory species.

### Seasonal warming sensitivity suggests temporally- and species-specific risk

Warming sensitivity, expressed as the slope of cold hardiness change per degree of warming for each collection, revealed clear differences between plant functional groups and phase structure across years (**Fig. 3**). Here we explored sensitivity as a cumulative intra-seasonal temperature effect, quantified as a response of cold hardiness to the cumulative mean temperature difference between each plot and ambient reference plots from 1 August through each collection date (ΔT) within each season. Under these conditions, sensitivity exceeding 1°C per °C of warming indicates that cold hardiness loss outpaces the change in temperature by warming treatments, thus elevating cold damage risk by effectively reducing safety margins, whereas sensitivities below 1 mean lower cold damage risk with warming.

**Figure 3.**
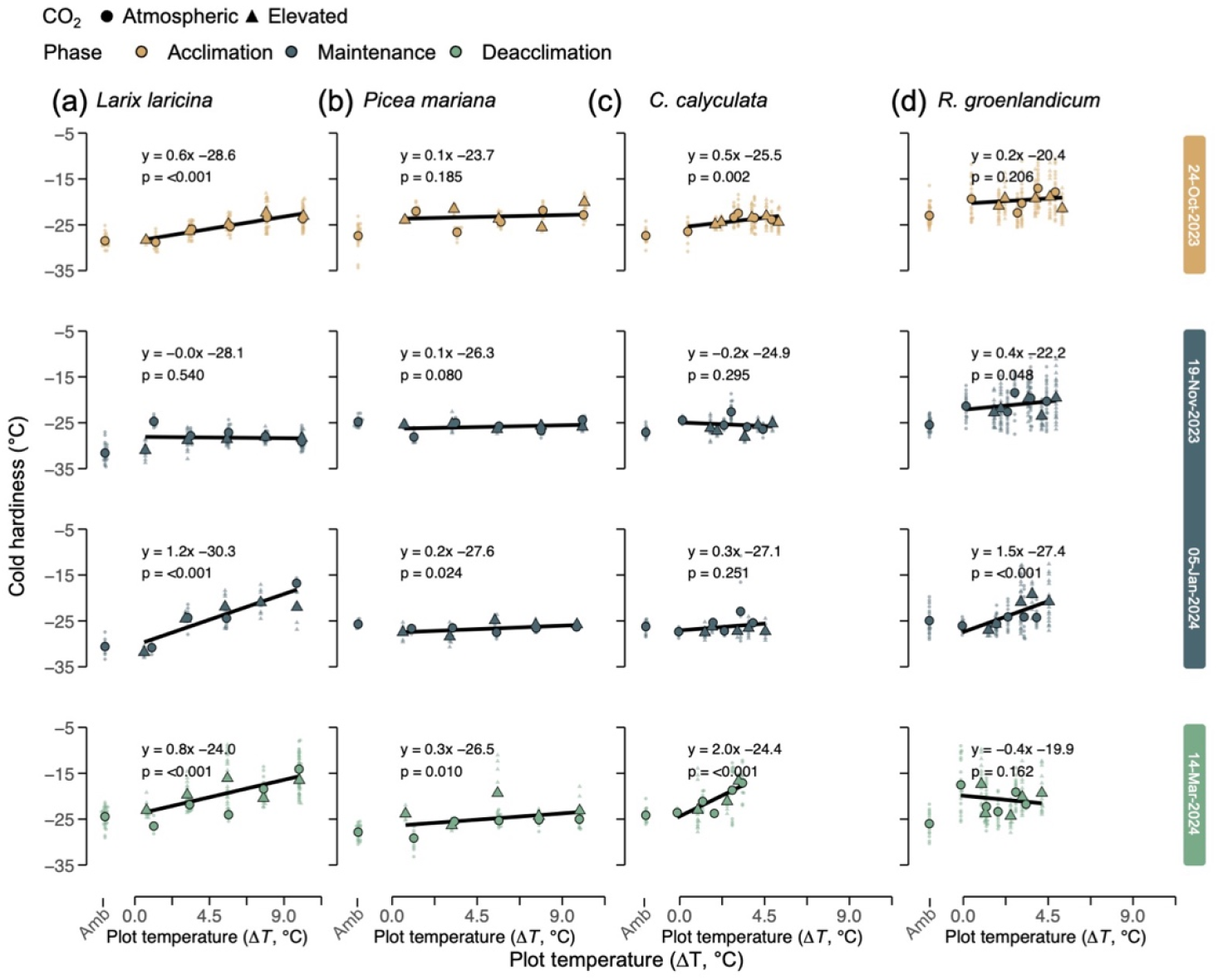
Phase-specific cold hardiness responses to warming. Cold hardiness measured at each warming level during the 2023–2024 season for *Larix laricina* (**a**), *Picea mariana* (**b**), *Chamaedaphne calyculata* (**c**) and *Rhododendron groenlandicum* (**d**), organized by seasonal phase: acclimation (golden), maintenance (dark teal), and deacclimation (sage green). Small points show single bud measurements, while larger points show averages within a plot for each collection date. Solid lines represent linear regression fits. Ambient plots are the reference for temperature (i.e. a ΔT of 0 means temperature equal to ambient plots). Full seasonal trajectories for all species and years are shown in **Fig. S3**; additional treatment- and collection-level patterns are shown in **Figs. S9–S12**.

The cumulative effect of temperature differed for the two functional groups. For overstory species, warming treatments produced relatively even warming throughout the dormant season, with ΔT minimum and maximum ranges of 8.54°C to 9.25°C between the +0.00°C and +9.00°C treatments (i.e. distribution of points along the x-axis within **Fig. 3a,b**, and y-axis within **Fig. S5**). For understory species, however, snow cover fluctuations resulted in more variable conditions, including temperature inversions related to target treatments. This resulted in a compressing of the effective ΔT range over time (2.02°C to 4.50°C; **Fig. 3c,d, S5**).

The calculation of sensitivity is exemplified for four timepoints (**Fig. 3**). The effect of ΔT on cold hardiness is significant during acclimation but shifts to non-significant during the maintenance period. An exceptionally warm period in late December 2023 and early January 2024 (with temperatures ∼7.0°C above seasonal norms for December (**Fig. S6**) and largely above freezing for 17 days in the warmest plot) triggered transient midwinter deacclimation in warmed plots, leading to high sensitivity in all species with the exception of leatherleaf (**Fig. 3**). During spring deacclimation, warming sensitivity returned to significant levels.

When explored over time, all four species show similarities in the trajectories of sensitivity (**Fig. 4**). In August, sensitivity is near zero as buds have only their inherent levels of cold hardiness as they are formed regardless of summer temperatures. Slight increases in sensitivity are observed during acclimation and reduced during maintenance as plants reach their maximum cold hardiness. All species show strong increases in sensitivity during the deacclimation phase. These seasonal sensitivity patterns were similar when inter-collection mean temperature (our measurement of short-term temperature responses) is used as the predictor instead of ΔT (**Fig. S7**), suggesting that the observed dynamics are robust and not dependent on the temperature metric used.

**Figure 4.**
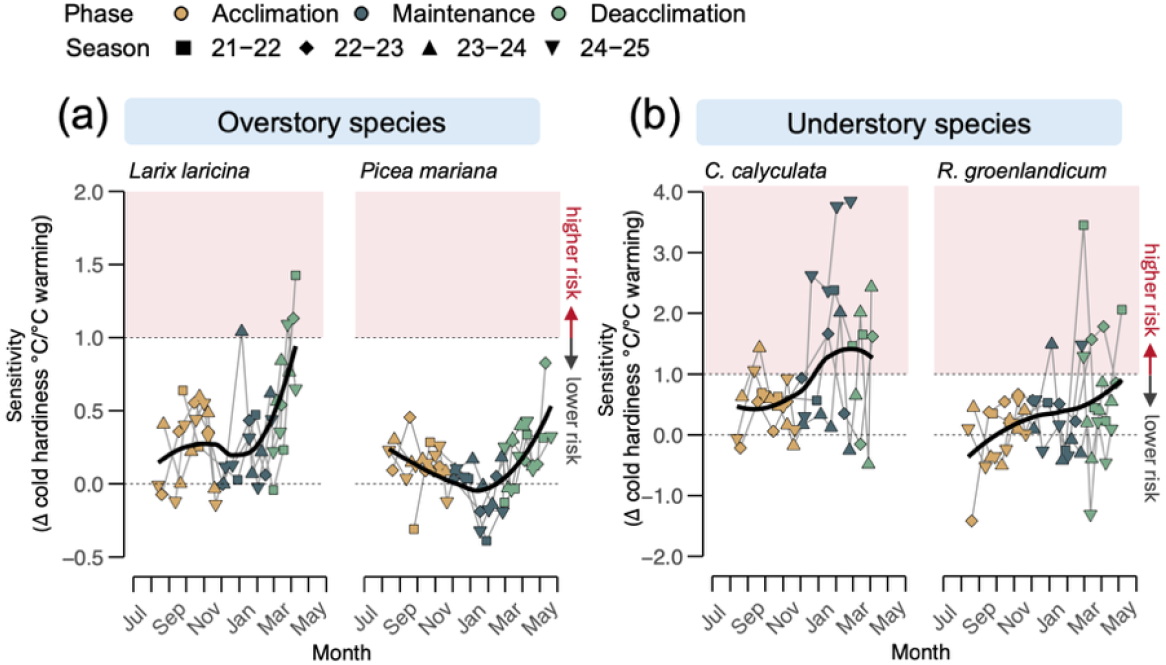
Warming sensitivity of bud cold hardiness over the dormant season. Slopes of bud cold hardiness in response to cumulative warming changes in temperature experienced by plants for (**a**) overstory and (**b**) understory species across acclimation (golden), maintenance (dark teal), and deacclimation (sage green) phases. Each point represents the slope of cold hardiness change per °C warming using ΔT for an individual collection across four seasons. Thick black lines show the loess-smoothed trend; thin lines connect points within individual seasons. A sensitivity > 1°C/°C indicates higher cold damage risk or reduced safety margins, whereas sensitivity < 1°C/°C indicates lower risk. A sensitivity = 0°C/°C means no effect of temperature. Only slopes with n ≥ 4 treatments were retained for overstory and understory species.

Among overstory species, larch was more sensitive to warming than black spruce across all phases (**Fig. 4a**). At the end of an unusual midwinter warm spell spanning late 2023 and early 2024, larch showed a sensitivity of ∼1.2°C per °C on 5 January 2024 (**Fig. 3a**), while black spruce exhibited a weaker response (∼0.2°C per °C; **Fig. 3b**). Both species fully reacclimated when cold temperatures returned. Importantly, at the end of the deacclimation period, larch showed sensitivity levels closer to and above 1.0°C per °C, whereas black spruce still showed comparatively more conservative responses even during this most responsive phase (average of ∼0.5°C per °C).

Understory species showed similarly phase-dependent responses (**Fig. 3c,d, 4b**), but their magnitude was strongly modulated by warming-induced reductions in snow cover. This resulted in sensitivities higher than 1°C per °C occurring earlier in the season, and to much greater magnitudes in both directions compared to overstory species. Understory species also showed higher year to year variation that reflect different snow conditions across years.

Changes in cold damage risk were not uniform across the dormant season. While cold hardiness was most often significantly affected by treatments during acclimation (sensitivity > 0°C per °C), cold damage risk was always reduced with increasing temperatures in this phase (sensitivity ≤1°C per °C). During the maintenance period, risk was substantially reduced with warming for most species. Higher risk of bud cold damage with warming (sensitivity > 1°C per °C) was observed in larch and understory species with progression of the dormant season, but not in black spruce. However, for the three species that crossed that threshold, the time when it occurred within the season varied. Highest risk was observed in understory species, amplified by snow cover loss that disproportionately increased their exposure to temperature extremes, resulting in observable bud damage in warmer plots (**Fig. S8**).

## Discussion

Here we examined the effects of experimental whole-ecosystem warming on cold hardiness dynamics in boreal woody perennial plant species. Our results demonstrate that climate warming alters cold hardiness dynamics in both overstory and understory species, but the magnitude and consequences of these responses vary between seasonal phases and species. Despite widespread warming across northern forests over the past century (*31, 32*), we found no evidence that warming alters maximum midwinter cold tolerance – even at the warm range edge of the boreal forest where this experiment is located. Instead, warming delayed fall acclimation and most importantly accelerated spring deacclimation, particularly increasing plant vulnerability during the spring transitional period when cold hardiness declines faster than environmental cold risk recedes. These phase- and species-specific responses reduce safety margins most strongly during transitional seasons and suggest that climate-driven shifts in seasonal timing may increasingly influence cold damage risk and future species composition in boreal forests.

Elevated CO_2_ did not influence cold hardiness dynamics across species or phases. Cold hardiness is related to carbohydrate concentrations, including the cold hardiness of buds as they form in the summer (*33*). CO_2_ enrichment has been shown to not increase concentration of non-structural carbohydrates in tissues, even when total growth increases (*34*). Our results suggest that current CO_2_ levels pose no direct limitation on carbon metabolism related to cold hardiness – even at higher temperatures where respiration can be higher (*35*). Though rising atmospheric CO_2_ concentration will not directly alter the freeze risk of perennial plants, its effect in natural ecosystems will be a result of its greenhouse gas properties increasing temperature.

Incremental warming delayed fall acclimation and accelerated spring deacclimation, together extending the period during which tissues were most susceptible to damaging temperatures at both ends of the dormant season (**Fig. 1**). The delay in fall acclimation reflects a temperature-driven shift in the timing of cold hardiness acquisition rather than a reduction in the inherent capacity of boreal species to withstand extreme winter temperatures (*5, 6, 36*). Cold hardiness gains still occur at a fast pace despite warm temperatures (**Fig. 2**), leaving wide safety margins during fall. In spring, warming sensitivity was strongest, substantially narrowing the safety margin against late frost damage for larch and understory species. This phenological mismatch where the cold hardiness loss outpaces the seasonal increase in temperature is increasingly recognized as a key driver of cold damage in forests and woody perennial crops (*37*–*39*).

The exceptional warm period in December 2023 and January 2024 illustrates that increased cold damage risk derived from warming is not restricted to the spring deacclimation phase. Warming events that are intense and long enough, particularly occurring late during winter or early spring, can cause deacclimation that can be detrimental if reacclimation does not occur fast enough to avoid damage from returning intense cold (*19*), which here we observe as highly species-specific effects. Rather than altering the maximum depth of midwinter hardiness even at the warm range edge of the boreal forest, warming effects were expressed almost exclusively during seasons when plants were actively gaining or losing cold hardiness, highlighting that studies using only midwinter measurements may substantially underestimate climate-driven shifts in vulnerability.

Seasonal sensitivity shifts of cold hardiness responses reflect the physiological regulation of dormancy. During winter maintenance, buds remain dormant and exhibit limited responsiveness to temperature variation (**Figs. 3**,**4**), consistent with physiological constraints that stabilize cold hardiness against short-term fluctuations (*11, 20*). In contrast, plants become more temperature-responsive with dormancy progression (*12, 19, 40, 41*) and even brief warm periods during these phases can rapidly accelerate cold hardiness loss while damaging freeze events remain likely. Short-term temperature analyses confirmed this mechanism directly, showing that cold hardiness responsiveness was minimal during the maintenance phase, but higher during acclimation and deacclimation. Particularly, the high positive ΔCH even at low above-freezing temperatures during the deacclimation phase identifies late winter and spring as the period of greatest warming-driven freeze risk (**Fig. 2**).

Species differed strongly in their sensitivity to warming during transitional seasons, reflecting contrasting life-history strategies along the carbon acquisitive-conservative axis. As a deciduous conifer, larch exhibited rapid loss of cold hardiness as temperatures increased in spring (**Fig. 1**) and showed higher sensitivity during transitional phases compared to black spruce (**Fig. 4**). Higher sensitivity may be the result of an acquisitive strategy in which early phenological responsiveness maximizes the potential growing season but increases exposure to episodic cold events during transitions as cold hardiness loss in deciduous species precedes budbreak and the onset of photosynthesis (*11, 42, 43*). Black spruce maintained slower and more conservative responses across all phases. As an evergreen conifer, black spruce retains foliage and the capacity to resume photosynthesis soon after warm spring temperatures return – in fact showing higher sensitivity for spring “green-up” than larch (*38*), without requiring rapid bud deacclimation and budbreak to secure carbon gain (*42, 44, 45*). The more rapid foliage green-up with warming in black spruce resulted in frost damage to foliage in April 2016 at SPRUCE, but not to buds (*38*), suggesting different cold hardiness dynamics may exist for foliage and buds in evergreen species. These contrasting strategies in carbon acquisition for deciduous and evergreen species produced substantially different bud cold hardiness safety margins under warming. Therefore, both warming and greater temperature variability resulting from climate change may increasingly disadvantage the acquisitive species with higher cold hardiness sensitivity relative to more conservative counterparts, resulting in overstory composition shifts due to cold damage.

Warming also altered cold vulnerability through indirect microclimatic pathways that disproportionately affected understory species. Reduced snow cover in warmer enclosures eliminated the insulating buffer that normally moderates near-surface winter temperatures (*38, 46, 47*). Paradoxically, this exposed understory buds in the plots with the highest warming treatments to temperatures colder than within ambient and least warmed plots (**Fig. S1**), such that measured microclimate temperatures were not proportional to the intended warming treatment. This temperature reversal likely explains the non-significant and occasionally negative sensitivities observed during spring deacclimation, underscoring that surface-level microclimate and cold hardiness dynamics cannot be reliably inferred from typical weather station height air temperature in ecosystems where snow dynamics are changing (*29, 48*–*50*). Reduced snow cover observed in warmer plots in the SPRUCE experimental system may partially result from the methods used for warming and should be viewed as one plausible pattern with future real-world warming. That is, future snow depths in non-experimental systems may vary from those available for study in SPRUCE depending on changes in precipitation patterns, wind dynamics, and other factors not captured by the experiment.

The greater range of ΔCH in smaller range of temperatures observed in understory species during deacclimation (**Fig. 2b**) likely reflects their dependence on snow cover as a thermal buffer that typically moderates spring temperature exposure in boreal understories (*51, 52*). The distinct warming sensitivities observed between the two understory species, despite identical enclosure conditions, further suggest that species-specific tolerance to microclimatic variability, rather than mean warming alone, will determine understory compositional change under future climate scenarios (*50, 52*–*55*). Critically, bud damage was observed in both understory species in the warming treatments – particularly +6.75 and +9.00ºC – during two of the four study seasons: driven by a warm spell in December 2023 and early January 2024 during the 2023-24 season, and by warming-induced snow cover loss during the 2024-25 season (**Fig. S8**). This is consistent with longstanding experimental evidence that reduced snow insulation exposes understory plants to damaging freeze events (*29, 48, 49*). Here we provide experimental evidence that near-term air warming will cause cold injury through midwinter snowmelt in boreal plant communities.

Together, our results demonstrate that cold hardiness is a dynamic, phase-related trait, and that species-specific representation of these dynamics, which is limited in the literature (*12, 56*–*58*), is essential for predicting forest resilience, productivity, and climate feedbacks under warming. Studies of species distribution or productivity that assume constant cold hardiness or neglect phase-specific sensitivity risk may thus systematically under or overestimate climate-driven damage as winter and spring temperature variability intensifies, depending on when sensitivity is evaluated (*19, 36, 38*). Although warming may lengthen growing seasons for some species, increased exposure to episodic cold events during transitional seasons, especially in spring and amplified by snow cover loss in understory microclimates, may offset these gains through tissue damage or shifts in competitive dynamics. For species of low stature, snow loss under warming was associated with bud mortality in understory shrubs, highlighting snow cover as a main factor of winter survival. Differences in seasonal cold hardiness responses among species with contrasting carbon acquisition strategies may alter competitive advantages and facilitate range shifts under continued climate warming, particularly among species with more acquisitive growth habits (*13, 59, 60*). Integrating temporally determined vulnerability, microclimatic heterogeneity, and snow-mediated insulation into predictive models will therefore be critical for accurately projecting future changes in carbon sequestration, biodiversity, and ecosystem services across temperate and boreal forests.

## Materials and methods

### Site description and experimental design

Plant material was collected from trees and shrubs within the SPRUCE (Spruce and Peatland Responses Under Changing Environments) experiment. SPRUCE is located at the Marcell Experimental Forest in Minnesota, USA, at the S1-Bog, in the southern edge of the boreal zone (47° 30.476′□N; 93° 27.162′□W) (**Fig. 5a**). The S1-Bog is an ombrotrophic peatland with large and rapid diurnal and seasonal temperature fluctuations (subhumid continental climate) (*61*). The mean annual temperature, based on a 48-year average (1961-2009) is 3.4ºC, with recorded air temperatures ranging from –45ºC to 38ºC throughout the year (*31*). Canopy vegetation is dominated by the tree species black spruce [*Picea Mariana* (Mill.) B.S.P] and American larch [*Larix laricina* (Du Roi) K. Koch]. The understory includes ericaceous shrubs, from which the dominant two species were included in collections: Labrador tea [*Rhododendron groenlandicum* (Oeder) Kron & Judd], and leatherleaf [*Chamaedaphne calyculata* (L.) Moench.] (*30*)

**Figure 5.**
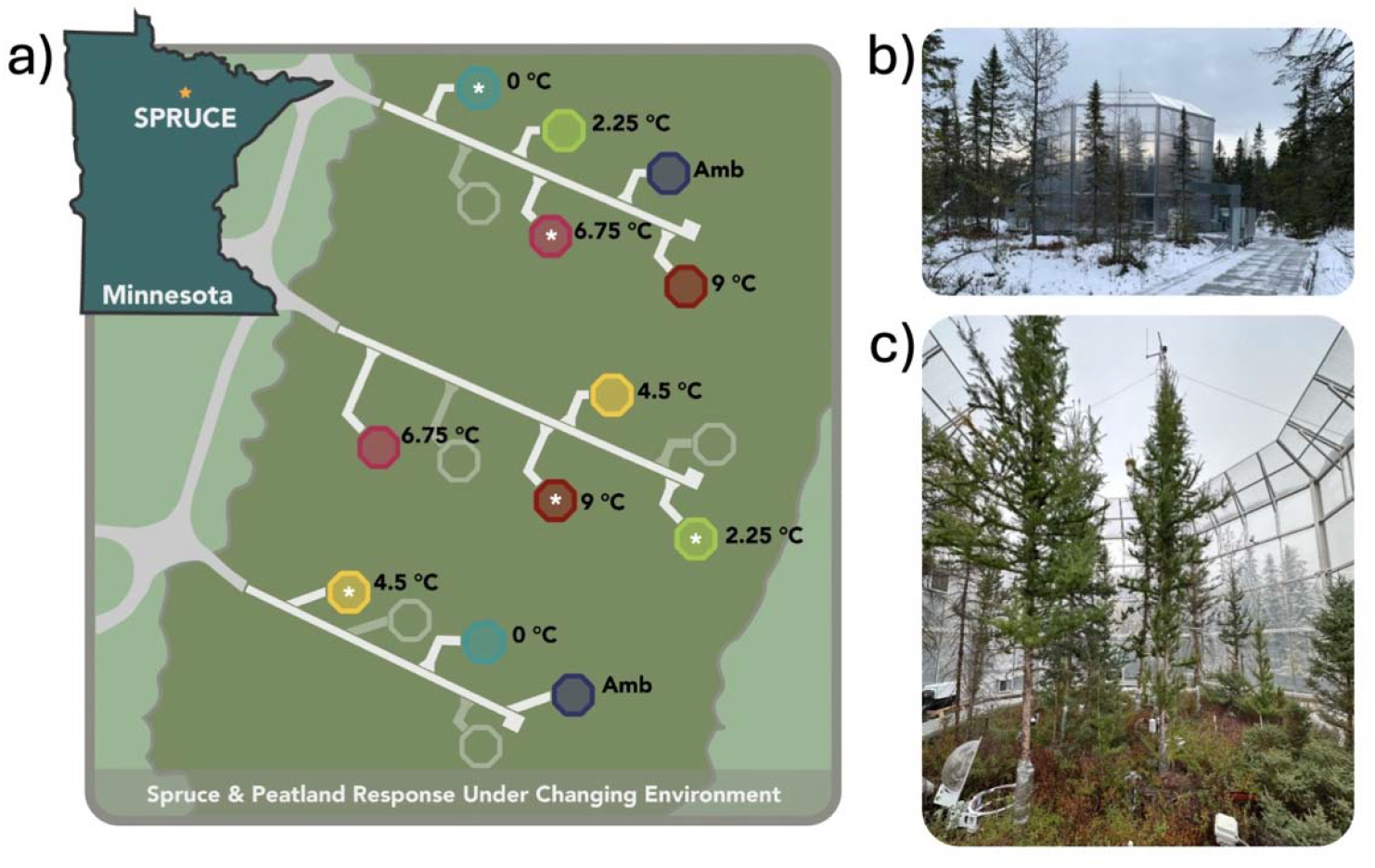
Layout of the SPRUCE experimental enclosures and associated treatments. (**a**) Location of the SPRUCE experimental site within the Marcell Experimental Forest in northern Minnesota, and schematic of the arrangement of enclosures and their assigned warming treatments, ranging from +0°C to +9□°C above ambient. Plots with an asterisk (*) were enriched with CO_2_ at ∼500ppm above atmospheric. (**b**) Exterior view of an experimental enclosure. (**c**) Interior view of an experimental enclosure.

Climate conditions were manipulated within ten large open-top enclosures (12.8m diameter, 7m height) built surrounding plants in 2015 (**Fig. 5b,c**). Each enclosure was constructed with double-walled transparent greenhouse panels to allow increase in both air and soil temperature, and in CO_2_ concentration. Additionally, a subsurface corral provides hydrologic isolation, allowing the soil to be warmed to match levels of air temperature warming (*30*).

Five levels of continuous ecosystem warming were applied, with target temperature treatment increases of: +0.00ºC, +2.25ºC, +4.50ºC, +6.75ºC, and +9.00ºC above non-warmed conditions. Experimental warming conditions were calibrated based on a non-heated control plot (+0.00 ºC), and also compared to temperature monitored in two adjacent non-enclosed plots, here referred to as “Amb” or “Ambient”. Temperatures were recorded at various aboveground heights and belowground depths at 30-min intervals across each plot. For these analyses, we used air temperature measured at 2 m height for overstory species and temperature at the soil surface (0 m) for understory shrubs. Two enclosures were assigned to each warming level, with one enclosure per level also receiving elevated CO_2_ (∼500 ppm above atmospheric). The SPRUCE project operated year-round with deep-soil heating initiated in June 2014, air warming beginning in August 2015 (*62*) and CO_2_ fumigation starting in June 2016 (*63*). The enclosures were constantly heated using propane-fired heat exchangers and a system of air blowers and conduits. A more detailed description of the experimental setup and treatments is provided within (*30*).

### Cold hardiness determination

Bud cold hardiness was monitored across four seasons (2021–2025) for all four species using differential thermal analysis (DTA) as described by (*64*). Briefly, buds were excised from the twigs and shoots avoiding damage to their internal structure and placed on thermoelectric modules (TEMs). Samples were then exposed to decreasing temperature in an environmental chamber. The temperature program held temperature at –5 °C for 5 hours, followed by a decrease to –55°C at a rate of –6°C h^−1^, and a final hold at –55°C for 2 hours. Low temperature exotherms (LTEs) were recorded using to a Keithley 6510 multimeter data acquisition system (Keithley Instruments), with LTE signals referenced to temperatures recorded by a thermocouple (22 AWG). DTA is an appropriate technique for species that use supercooling in their buds, which includes those studied here (*8*), and cold hardiness measurements were supported through a comparison to the visual browning method for cold hardiness determination (**Fig. S13**; (*65*)).

Samples were collected at semi-regular intervals from August through May of each of the four seasons. Buds were placed in plastic bags with moist tissue paper, transported in a cooler with ice and water mix (0 ºC) and stored at 4ºC until analysis. Occasionally, buds were shipped overnight in a cooler with ice packs, which does not significantly affect cold hardiness measurements (*66*). To minimize destructive sampling within plots, bud samples were pooled from multiple individuals of the same species within each enclosure. Typically, 10 to 15 buds were used to assess cold hardiness per species per enclosure (∼20,500 total buds for cold hardiness measurements over four years for all four species). Outliers were identified and removed using studentized residuals exceeding ±2.95 prior to analyses.

### Temperature characterization

Temperature effects on cold hardiness were examined at two temporal scales: inter-collection responses capturing the immediate effect of thermal conditions between consecutive sample collection dates (short-term), and responses reflecting the cumulative intra-seasonal thermal environment experienced by each plot from mid-Summer until the time of each sample collection within a season.

#### Inter-collection temperature effects

To characterize the thermal conditions experienced between consecutive sample collections, we calculated the mean air temperature over the inter-collection interval preceding each sampling date. For each treatment plot and season, the inter-collection (ic) mean temperature 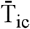for collection date *c* was computed as:

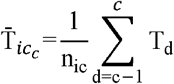

where T_d_ is the daily mean temperature on day d within the interval between collection date *c-1* and collection date *c*, and n_ic_ is the number of days in that inter-collection interval. This variable captures the thermal conditions most immediately preceding each cold hardiness measurement and was used to assess short-term responsiveness of cold hardiness to recent temperature in absolute values.

#### Cumulative intra-seasonal temperature effects

To characterize the cumulative thermal environment of each plot across the season, we computed the average relative temperature adjustment (effective ΔT) for each plot in relation to ambient temperature plots. For each collection date *c*, the cumulative mean temperature for each plot 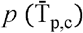 was calculated as the mean of all daily temperatures from 1 August of through each collection date *c* within a season:

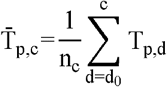

where *d*_0_ is 1 August and n_c_ is the number of days from *d*_0_ to collection date *c*. The reference temperature for each collection was defined as the mean of 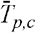 across the two ambient reference plots (both non-warmed and without enclosure). The effective temperature differential between any plot and ambient plot temperatures (ΔT) was then calculated as:

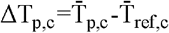

where 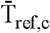 is the cumulative mean temperature of the reference plots for each collection within a season. Thus, ΔT represents how much warmer each plot was relative to ambient conditions up to each sampling point and was used as the predictor of cumulative intra-seasonal warming effects on cold hardiness.

### Data analysis and statistical tests

All data were analyzed using R [ver. 4.4.0; (*67*)] within R Studio (*68*). Given the regression-based experimental design, warming treatment was treated as a continuous variable throughout, represented by ΔT or 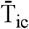, and only first-degree polynomial regressions were considered.

#### CO_2_ and treatment effects

A full model was fit for cold hardiness using CO_2_ concentration, plot temperature ΔT, and collection as explanatory variables. Cold hardiness was not significantly affected by elevated CO_2_ or its interaction with warming treatment across species (**Notes S1, Table S1**), and CO_2_ was therefore excluded from all subsequent analyses.

#### Short-term thermal responsiveness

To quantify short-term sensitivity of cold hardiness to recent temperature conditions, we calculated the change in cold hardiness between consecutive sampling dates as:

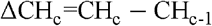

where CH_c_ is the mean cold hardiness on collection date *c* and CH_c-1_ is the mean on the previous sampling date, calculated separately for each species, treatment, and season. ΔCH values were then matched to their corresponding inter-collection mean temperatures 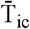and each observation was assigned to its respective seasonal phase. To characterize phase-specific thermal responsiveness, phase centroids were calculated as the mean inter-collection temperature and mean ΔCH for all observations within each phase and functional group. Bivariate 95% confidence ellipses were fitted assuming a normal distribution using stat_ellipse() in ggplot2 (*69*) to visualize the dispersion of observations around each centroid and assess overlap between phases.

#### Cumulative intra-seasonal warming sensitivity

To quantify the sensitivity of cold hardiness to cumulative intra-seasonal warming at each collection, separate linear models were fitted for each species and collection using ΔT as the sole predictor:

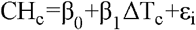

where CH_c_ is cold hardiness for collection *c*, β_0_ is the intercept representing cold hardiness at approximately ambient conditions (ΔT = 0°C), β_1_ is the slope associated with the cold hardiness sensitivity to warming (°C cold hardiness change per °C of warming), and ε_i_ is the residual error. Slopes were estimated only for collection periods with at least four unique target temperature levels represented to ensure stable parameter estimation. Slopes were estimated separately for each collection to capture phase-specific variation in warming sensitivity across the season.

To account for seasonal variation in the magnitude of both inter-collection and cumulative intra-seasonal temperature responses, data were grouped into three phases based on overall cold hardiness patterns observed across species and years for interpretation: acclimation (1 Aug – 31 Oct), maintenance (1 Nov – 15 Mar), and deacclimation (16 Mar – 30 Jun).

## Acknowledgments

We thank J. J. Grossman, K. A. McCulloh and J. P. Londo for comments on the work. This work is partially supported by the National Institute of Food and Agriculture, United States Department of Agriculture, McIntire Stennis projects 7002619 and 1027327, and by the Office of the Vice Chancellor for Research at the University of Wisconsin–Madison with funding from the Wisconsin Alumni Research Foundation. This material is based upon work supported by the U.S. Department of Energy, Office of Science, Office of Biological and Environmental Research. Oak Ridge National Laboratory is managed by UT-Battelle, LLC, for the U.S. Department of Energy under contract DE-AC05-00OR22725. The SPRUCE experiment is a collaborative research effort between ORNL and the USDA Forest Service.

## Author contributions

Conceptualization: APK, PJH; Data Curation: FC-A, APK; Formal Analysis: FC-A, APK; Funding Acquisition: APK, PJH; Investigation: FC-A, EK, MGN, APK; Resources: FC-A, EK, MGN, APK, KJP, MPG; Supervision: APK; Visualization: FC-A, APK; Writing – original draft: FC-A; Writing – review & editing: APK, MGN, EK, KJP, MPG, PJH.

## Supplementary Materials

Supplementary Text

Figs. S1 to S13

Table S1

## References

1. J. Parker, Cold Resistance in Woody Plants. Botanical Review 29, 123–201 (1963).

2. G. Neuner, K. Monitzer, D. Kaplenig, J. Ingruber, Frost Survival Mechanism of Vegetative Buds in Temperate Trees: Deep Supercooling and Extraorgan Freezing vs. Ice Tolerance. Front. Plant Sci. Volume 10-2019 (2019).

3. S. Kellomäki, H. Peltola, T. Nuutinen, K. T. Korhonen, H. Strandman, Sensitivity of managed boreal forests in Finland to climate change, with implications for adaptive management. Philosophical Transactions of the Royal Society B: Biological Sciences 363, 2339–2349 (2008).

4. H. Hänninen, “Climatic Adaptation of Boreal and Temperate Tree Species” in Boreal and Temperate Trees in a Changing Climate: Modelling the Ecophysiology of Seasonality, H. Hänninen, Ed. (Springer Netherlands, Dordrecht, 2016; 10.1007/978-94-017-7549-6_1), pp. 1–13.

5. F. S. Chapin, A. D. McGuire, R. W. Ruess, T. N. Hollingsworth, M. C. Mack, J. F. Johnstone, E. S. Kasischke, E. S. Euskirchen, J. B. Jones, M. T. Jorgenson, K. Kielland, G. P. Kofinas, M. R. Turetsky, J. Yarie, A. H. Lloyd, D. L. Taylor, Resilience of Alaska’s boreal forest to climatic change. Canadian Journal of Forest Research 40, 1360–1370 (2010).

6. D. Stralberg, D. Arseneault, J. L. Baltzer, Q. E. Barber, E. M. Bayne, Y. Boulanger, C. D. Brown, H. A. Cooke, K. Devito, J. Edwards, C. A. Estevo, N. Flynn, L. E. Frelich, E. H. Hogg, M. Johnston, T. Logan, S. M. Matsuoka, P. Moore, T. L. Morelli, J. L. Morissette, E. A. Nelson, H. Nenzén, S. E. Nielsen, M.-A. Parisien, J. H. Pedlar, D. T. Price, F. K. A. Schmiegelow, S. M. Slattery, O. Sonnentag, D. K. Thompson, E. Whitman, Climate-change refugia in boreal North America: what, where, and for how long? Front. Ecol. Environ. 18, 261–270 (2020).

7. A. Sakai, W. Larcher, “Regional Distribution of Plants and Their Adaptive Responses to Low Temperatures” in Frost Survival of Plants: Responses and Adaptation to Freezing Stress, A. Sakai, W. Larcher, Eds. (Springer Berlin Heidelberg, Berlin, Heidelberg, 1987; 10.1007/978-3-642-71745-1_7), pp. 174–234.

8. A. Sakai, W. Larcher, “Mechanisms of Frost Survival” in Frost Survival of Plants: Responses and Adaptation to Freezing Stress, A. Sakai, W. Larcher, Eds. (Springer Berlin Heidelberg, Berlin, Heidelberg, 1987; 10.1007/978-3-642-71745-1_4), pp. 59–96.

9. G. R. Strimbeck, P. G. Schaberg, C. G. Fossdal, W. P. Schröder, T. D. Kjellsen, Extreme low temperature tolerance in woody plants. Front. Plant Sci. Volume 6-2015 (2015).

10. J. Alden, R. K. Hermann, Aspects of the cold-hardiness mechanism in plants. The Botanical Review 37, 37–142 (1971).

11. Y. Vitasse, A. Lenz, C. Körner, The interaction between freezing tolerance and phenology in temperate deciduous trees. Front. Plant Sci. Volume 5-2014 (2014).

12. A. P. Kovaleski, Woody species do not differ in dormancy progression: Differences in time to budbreak due to forcing and cold hardiness. Proceedings of the National Academy of Sciences 119, e2112250119 (2022).

13. J. P. Londo, A. P. Kovaleski, Integrating cold hardiness and deacclimation resistance demonstrates a conserved response to chilling accumulation in grapevines. J. Exp. Bot., eraf045 (2025).

14. F. Campos-Arguedas, E. Kirchhof, M. G. North, J. P. Londo, T. Bates, C. van Leeuwen, A. Destrac-Irvine, B. Bois, A. P. Kovaleski, Cold hardiness dynamics predict budbreak and associated low-temperature threats in grapevine. New Phytologist n/a (2026).

15. E. M. Wolkovich, C. J. Chamberlain, D. M. Buonaiuto, A. K. Ettinger, I. Morales-Castilla, Integrating experiments to predict interactive cue effects on spring phenology with warming. New Phytologist 235, 1719–1728 (2022).

16. A. K. Ettinger, C. J. Chamberlain, I. Morales-Castilla, D. M. Buonaiuto, D. F. B. Flynn, T. Savas, J. A. Samaha, E. M. Wolkovich, Winter temperatures predominate in spring phenological responses to warming. Nat. Clim. Chang. 10, 1137–1142 (2020).

17. G. Neuner, B. Kreische, D. Kaplenig, K. Monitzer, R. Miller, Deep supercooling enabled by surface impregnation with lipophilic substances explains the survival of overwintering buds at extreme freezing. Plant Cell Environ. 42, 2065–2074 (2019).

18. P. J. CaraDonna, J. A. Bain, Frost sensitivity of leaves and flowers of subalpine plants is related to tissue type and phenology. Journal of Ecology 104, 55–64 (2016).

19. A. P. Kovaleski, The potential for an increasing threat of unseasonal temperature cycles to dormant plants. New Phytologist 244, 377–383 (2024).

20. A. Lenz, G. Hoch, C. Körner, Y. Vitasse, Convergence of leaf-out towards minimum risk of freezing damage in temperate trees. Funct. Ecol. 30, 1480–1490 (2016).

21. J. Wang, H. Hua, J. Guo, X. Huang, X. Zhang, Y. Yang, D. Wang, X. Guo, R. Zhang, N. G. Smith, S. Rossi, J. Peñuelas, P. Ciais, C. Wu, L. Chen, Late spring frost delays tree spring phenology by reducing photosynthetic productivity. Nat. Clim. Chang. 15, 201–209 (2025).

22. C. J. Chamberlain, B. I. Cook, I. García de Cortázar-Atauri, E. M. Wolkovich, Rethinking false spring risk. Glob. Chang. Biol. 25, 2209–2220 (2019).

23. D. R. Weisz, E. D. Skillman, J. M. Cannon al -, M.-X. Zhao, G.-M. Le, Y.-H. Liu -, C. Li, L. Wang, G. P. Marino, D. P. Kaiser, L. Gu, D. M. Ricciuto, Reconstruction of false spring occurrences over the southeastern United States, 1901–2007:an increasing risk of spring freeze damage? Environmental Research Letters 6, 024015 (2011).

24. J. R. Lamichhane, Rising risks of late-spring frosts in a changing climate. Nat. Clim. Chang. 11, 554–555 (2021).

25. L. Maracahipes, M. B. Carlucci, E. Lenza, B. S. Marimon, B. H. Marimon, F. A. G. Guimarães, M. V Cianciaruso, How to live in contrasting habitats? Acquisitive and conservative strategies emerge at inter-and intraspecific levels in savanna and forest woody plants. Perspect. Plant Ecol. Evol. Syst. 34, 17–25 (2018).

26. J. Luo, J. Li, Y. Sun, A. A. Degen, X. Wang, J. Xiong, M. Aqeel, S. Xie, D. Tang, F. Li, W. Hu, L. Dong, J. Peng, Q. Hou, Y. Deng, J. Ran, J. Deng, Differential resource acquisition strategies of herbaceous and woody plants in temperate forests. BMC Plant Biol. 26, 196 (2026).

27. L. Gu, P. J. Hanson, W. Mac Post, D. P. Kaiser, B. Yang, R. Nemani, S. G. Pallardy, T. Meyers, The 2007 Eastern US Spring Freeze: Increased Cold Damage in a Warming World? Bioscience 58, 253–262 (2008).

28. P. G. Schaberg, D. V D’Amore, P. E. Hennon, J. M. Halman, G. J. Hawley, Do limited cold tolerance and shallow depth of roots contribute to yellow-cedar decline? For. Ecol. Manage. 262, 2142–2150 (2011).

29. J. Kreyling, M. Haei, H. Laudon, Absence of snow cover reduces understory plant cover and alters plant community composition in boreal forests. Oecologia 168, 577–587 (2012).

30. P. J. Hanson, J. S. Riggs, W. R. Nettles, J. R. Phillips, M. B. Krassovski, L. A. Hook, L. Gu, A. D. Richardson, D. M. Aubrecht, D. M. Ricciuto, J. M. Warren, C. Barbier, Attaining whole-ecosystem warming using air and deep-soil heating methods with an elevated CO2 atmosphere. Biogeosciences 14, 861–883 (2017).

31. S. D. Sebestyen, C. Dorrance, D. M. Olson, E. S. Verry, R. K. Kolka, A. E. Elling, R. Kyllander, “Long-term monitoring sites and trends at the Marcell Experimental Forest” in Peatland Biogeochemistry and Watershed Hydrology at the Marcell Experimental Forest (CRC press Boca Raton, FL, 2011) vol. 1, pp. 15–71.

32. A. R. Contosta, N. J. Casson, S. Garlick, S. J. Nelson, M. P. Ayres, E. A. Burakowski, J. Campbell, I. Creed, C. Eimers, C. Evans, I. Fernandez, C. Fuss, T. Huntington, K. Patel, R. Sanders-DeMott, K. Son, P. Templer, C. Thornbrugh, Northern forest winters have lost cold, snowy conditions that are important for ecosystems and human communities. Ecological Applications 29, e01974 (2019).

33. X. Morin, T. Améglio, R. Ahas, C. Kurz-Besson, V. Lanta, F. Lebourgeois, F. Miglietta, I. Chuine, Variation in cold hardiness and carbohydrate concentration from dormancy induction to bud burst among provenances of three European oak species. Tree Physiol. 27, 817–825 (2007).

34. R. Tognetti, J. Johnson D., The effect of elevated atmospheric CO2 concentration and nutrient supply on gas exchange, carbohydrates and foliar phenolic concentration in live oak (Quercus virginiana Mill.) seedlings. Ann. For. Sci. 56, 379–389 (1999).

35. Y. Velappan, T. G. Chabikwa, J. A. Considine, P. Agudelo-Romero, C. H. Foyer, S. Signorelli, M. J. Considine, The bud dormancy disconnect: latent buds of grapevine are dormant during summer despite a high metabolic rate. J. Exp. Bot. 73, 2061–2076 (2022).

36. G. Neuner, Frost resistance in alpine woody plants. Front. Plant Sci. Volume 5-2014 (2014).

37. C. Bigler, H. Bugmann, Climate-induced shifts in leaf unfolding and frost risk of European trees and shrubs. Sci. Rep. 8, 9865 (2018).

38. A. D. Richardson, K. Hufkens, T. Milliman, D. M. Aubrecht, M. E. Furze, B. Seyednasrollah, M. B. Krassovski, J. M. Latimer, W. R. Nettles, R. R. Heiderman, J. M. Warren, P. J. Hanson, Ecosystem warming extends vegetation activity but heightens vulnerability to cold temperatures. Nature 560, 368–371 (2018).

39. C. J. Chamberlain, E. M. Wolkovich, Late spring freezes coupled with warming winters alter temperate tree phenology and growth. New Phytologist 231, 987–995 (2021).

40. M. G. North, A. P. Kovaleski, Time to budbreak is not enough: cold hardiness evaluation is necessary in dormancy and spring phenology studies. Ann. Bot., doi: 10.1093/aob/mcad182 (2024).

41. M. G. North, B. A. Workmaster, A. Atucha, A. P. Kovaleski, Cold hardiness-informed budbreak reveals role of freezing temperatures and daily fluctuation in a chill accumulation model. J. Exp. Bot. 75, 6182–6193 (2024).

42. T. J. Givnish, Adaptive significance of evergreen vs. deciduous leaves: solving the triple paradox. Silva fennica 36, 703–743 (2002).

43. D. R. Bowling, C. Schädel, K. R. Smith, A. D. Richardson, M. Bahn, M. A. Arain, A. Varlagin, A. P. Ouimette, J. M. Frank, A. G. Barr, I. Mammarella, L. Šigut, V. Foord, S. P. Burns, L. Montagnani, M. E. Litvak, J. W. Munger, H. Ikawa, D. Y. Hollinger, P. D. Blanken, M. Ueyama, G. Matteucci, C. Bernhofer, G. Bohrer, H. Iwata, A. Ibrom, K. Pilegaard, D. L. Spittlehouse, H. Kobayashi, A. R. Desai, R. M. Staebler, T. A. Black, Phenology of Photosynthesis in Winter-Dormant Temperate and Boreal Forests: Long-Term Observations From Flux Towers and Quantitative Evaluation of Phenology Models. J. Geophys. Res. Biogeosci. 129, e2023JG007839 (2024).

44. M. E. Dusenge, J. M. Warren, P. B. Reich, E. J. Ward, B. K. Murphy, A. Stefanski, R. Bermudez, M. Cruz, D. A. McLennan, A. W. King, R. A. Montgomery, P. J. Hanson, D. A. Way, Boreal conifers maintain carbon uptake with warming despite failure to track optimal temperatures. Nat. Commun. 14, 4667 (2023).

45. G. Öquist, N. P. A. Huner, Photosynthesis of Overwintering Evergreen Plants. Annu. Rev. Plant Biol. 54, 329–355 (2003).

46. A. D. Richardson, T. F. Keenan, M. Migliavacca, Y. Ryu, O. Sonnentag, M. Toomey, Climate change, phenology, and phenological control of vegetation feedbacks to the climate system. Agric. For. Meteorol. 169, 156–173 (2013).

47. S. F. Bokhorst, J. W. Bjerke, H. Tømmervik, T. V Callaghan, G. K. Phoenix, Winter warming events damage sub-Arctic vegetation: consistent evidence from an experimental manipulation and a natural event. Journal of Ecology 97, 1408–1415 (2009).

48. F. C. Gates, The Relation of Snow Cover to Winter Killing in Chamaedaphne Calyculata. Torreya 12, 257–262 (1912).

49. G. Neuner, D. Ambach, K. Aichner, Impact of snow cover on photoinhibition and winter desiccation in evergreen Rhododendron ferrugineum leaves during subalpine winter. Tree Physiol. 19, 725–732 (1999).

50. P. De Frenne, F. Rodríguez-Sánchez, D. A. Coomes, L. Baeten, G. Verstraeten, M. Vellend, M. Bernhardt-Römermann, C. D. Brown, J. Brunet, J. Cornelis, G. M. Decocq, H. Dierschke, O. Eriksson, F. S. Gilliam, R. Hédl, T. Heinken, M. Hermy, P. Hommel, M. A. Jenkins, D. L. Kelly, K. J. Kirby, F. J. G. Mitchell, T. Naaf, M. Newman, G. Peterken, P. Petřík, J. Schultz, G. Sonnier, H. Van Calster, D. M. Waller, G.-R. Walther, P. S. White, K. D. Woods, M. Wulf, B. J. Graae, K. Verheyen, Microclimate moderates plant responses to macroclimate warming. Proceedings of the National Academy of Sciences 110, 18561–18565 (2013).

51. K. Charra-Vaskou, G. Charrier, A. Ganthaler, T. Améglio, S. Mayr, Reduced snow cover at the alpine treeline: resistance and recovery of saplings. New Phytologist 250, 1492–1509 (2026).

52. C. Steger, S. Kotlarski, T. Jonas, C. Schär, Alpine snow cover in a changing climate: a regional climate model perspective. Clim. Dyn. 41, 735–754 (2013).

53. T. Panchard, O. Broennimann, M. Gravey, G. Mariethoz, A. Guisan, Snow cover persistence as a useful predictor of alpine plant distributions. J. Biogeogr. 50, 1789–1802 (2023).

54. C. Marty, S. Schlögl, M. Bavay, M. Lehning, How much can we save? Impact of different emission scenarios on future snow cover in the Alps. Cryosphere 11, 517–529 (2017).

55. F. Zellweger, D. Coomes, J. Lenoir, L. Depauw, S. L. Maes, M. Wulf, K. J. Kirby, J. Brunet, M. Kopecký, F. Máliš, W. Schmidt, S. Heinrichs, J. den Ouden, B. Jaroszewicz, G. Buyse, F. Spicher, K. Verheyen, P. De Frenne, Seasonal drivers of understorey temperature buffering in temperate deciduous forests across Europe. Global Ecology and Biogeography 28, 1774–1786 (2019).

56. E. M. Wolkovich, B. I. Cook, J. M. Allen, T. M. Crimmins, J. L. Betancourt, S. E. Travers, S. Pau, J. Regetz, T. J. Davies, N. J. B. Kraft, T. R. Ault, K. Bolmgren, S. J. Mazer, G. J. McCabe, B. J. McGill, C. Parmesan, N. Salamin, M. D. Schwartz, E. E. Cleland, Warming experiments underpredict plant phenological responses to climate change. Nature 485, 494–497 (2012).

57. E. M. Wolkovich, B. I. Cook, T. J. Davies, Progress towards an interdisciplinary science of plant phenology: building predictions across space, time and species diversity. New Phytologist 201, 1156–1162 (2014).

58. F. A. M. Jones, C. Bogdanoff, E. M. Wolkovich, The role of genotypic and climatic variation at the range edge: A case study in winegrapes. Am. J. Bot. 111, e16270 (2024).

59. J. A. Savage, J. Cavender-Bares, Phenological cues drive an apparent trade-off between freezing tolerance and growth in the family Salicaceae. Ecology 94, 1708–1717 (2013).

60. A. Lenz, G. Hoch, Y. Vitasse, C. Körner, European deciduous trees exhibit similar safety margins against damage by spring freeze events along elevational gradients. New Phytologist 200, 1166–1175 (2013).

61. E. S. Verry, K. N. Brooks, P. K. Barten, “Streamflow response from an ombrotrophic mire” in In: Proceedings of the International Symposium on the Hydrology of Wetlands in Temperate and Cold Regions; 1988 June 6-8; Joensuu, Finland. Helsinki, Finland: International Peat Society/The Academy of Finland: 52–59. (1988).

62. P. J. Hanson, J. S. Riggs, W. R. Nettles, M. B. Krassovski, L. A. Hook, “SPRUCE whole ecosystems warming (WEW) environmental data beginning August 2015” (Oak Ridge National Laboratory (ORNL), Oak Ridge, TN (United States), 2016).

63. P. J. Hanson, W. R. Nettles, J. S. Riggs, M. B. Krassovski, L. A. Hook, “SPRUCE vertical profiles of CO2 and H2O concentrations in air in experimental plots beginning in 2015” (Oak Ridge National Lab.(ORNL), Oak Ridge, TN (United States), 2021).

64. L. J. Mills, J. C. Ferguson, M. Keller, Cold-hardiness evaluation of grapevine buds and cane tissues. Am. J. Enol. Vitic. 57, 194–200 (2006).

65. C. Villouta, B. A. Workmaster, J. Bolivar-Medina, S. Sinclair, A. Atucha, Freezing stress survival mechanisms in Vaccinium macrocarpon Ait. terminal buds. Tree Physiol. 40, 841–855 (2020).

66. J. P. Londo, M. M. Moyer, M. Mireles, L. Mills, M. Keller, B. A. Workmaster, A. Atucha, A. P. Kovaleski, Evaluation of Sample Preparation Practices Common with Differential Thermal Analysis of Grapevine Bud Cold Hardiness. Am. J. Enol. Vitic. 74, 0740002 (2023).

67. R Core Team, R: A Language and Environment for Statistical Computing. R Foundation for Statistical Computing [Preprint] (2024). https://www.R-project.org/.

68. Posit team, RStudio: Integrated Development Environment for R. Posit Software, PBC [Preprint] (2024). http://www.posit.co/.

69. H. Wickham, Elegant graphics for data analysis. Springer [Preprint] (2016).

