## Supplementary Materials for "Warming Reduces Cold Hardiness of Boreal Plants but Damage Risk Varies by Species and Season"

**The PDF file includes:**

Supplementary Text
Figs. S1 to S13
Table S1

**Supplementary Text**

**Notes S1. Model testing the effect of $CO_2$ on cold hardiness dynamics**

To assess the combined effects of warming treatment ($\Delta T$) and elevated $CO_2$ to cold hardiness ($CH_{ijk}$), we fitted a linear model using the *lm* function in R. The model included warming treatment ($\Delta T$), $CO_2$ concentration ($CO2_k$) and collection period ($Collection_j$) as fixed effects, along with interactions between all. The full model included:

$$CH_{ijk}=\beta_0+\beta_1\,\Delta T_{corrected_i}+\beta_2\,CO2_k+\beta_3\left(\Delta T_{corrected_i}\times CO2_k\right)+\beta_4\,Collection_j+$$
$$\beta_5\left(\Delta T_{corrected_i}\times Collection_j\right)+\beta_6\left(CO2_k\times Collection_j\right)+\beta_7\left(\Delta T_{corrected_i}\times CO2_k\times Collection_j\right)+\varepsilon_{ijk}$$

Here, $CH_{ijk}$ is the corrected cold hardiness for warming treatment i, $CO_2$ level $k$ and Collection j. $\beta_0$ is the overall intercept, while $\beta_1$, $\beta_2$ and $\beta_3$ are the fixed effects of warming treatment, $CO_2$, and their interaction, respectively. $\beta_4$ represents differences in baseline cold hardiness among collections; $\beta_5$, $\beta_6$, and $\beta_7$ represent collection-specific differences in Treatment and $CO_2$ sensitivity; and $\varepsilon_{ijk}$ is the residual error term.

Model simplification was performed by sequentially removing higher-order interaction terms and evaluating changes in model performance using Akaike's Information Criterion (AIC). Across species, $CO_2$ did not exhibit a consistent main effect on cold hardiness and terms involving $CO_2$ generally did not improve model performance relative to models accounting for collection and temperature responses (**Table

**S1**). An exception was observed for *C. calyculata*, where models including Treatment × $CO_2$ × Collection interactions showed marginally improved fit, suggesting weak, sequence-dependent modulation of temperature responses under elevated $CO_2$. However, these effects were small relative to the dominant influence of temperature and seasonal progression. Consequently, $CO_2$ as a factor was excluded from further analyses.

**Supplementary figures**

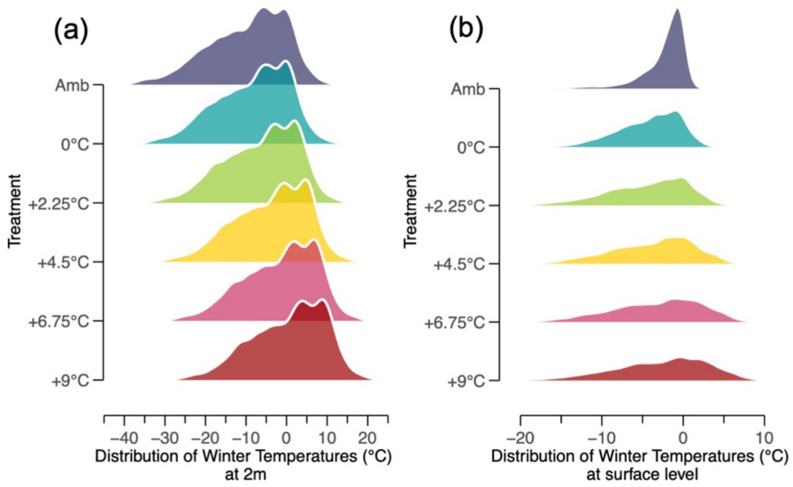


**Figure S1.** Histogram of measured winter temperatures at (**a**) 2 m and (**b**) surface level (0 m) over the course of four winter seasons at the SPRUCE experiment (1 August 2021 to 30 June 2025).

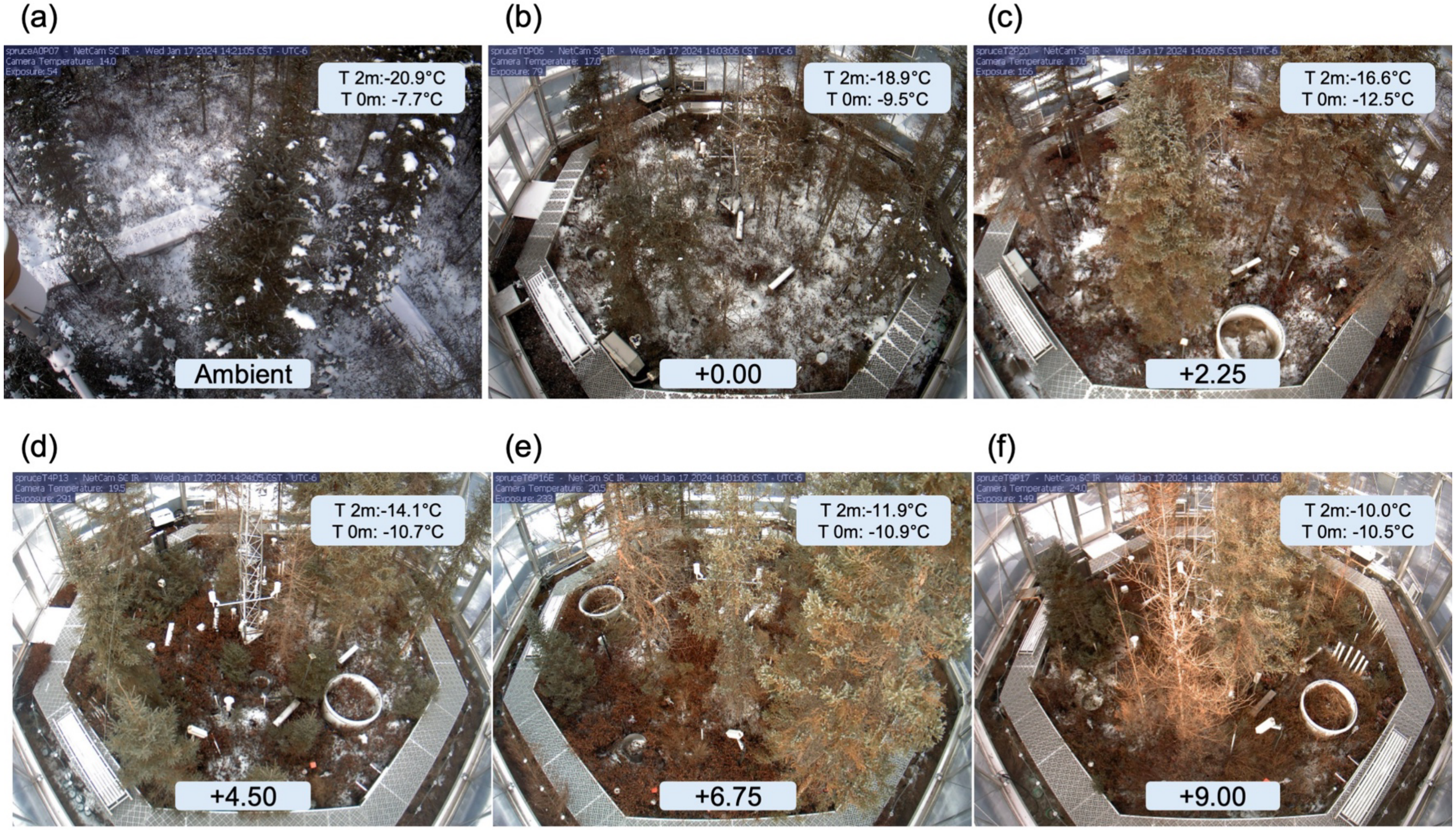


**Figure S2. Experimental warming reduces snow cover in SPRUCE enclosures.** Overhead images taken on the same day (January 17, 2024, ~2:00 PM) showing snow cover across warming treatments at the SPRUCE experiment. Panels show (a) ambient conditions and enclosures maintained at (b) +0.00°C, (c) +2.25°C, (d) +4.50°C, (e) +6.75°C, and (f) +9.00°C above ambient. Increasing warming levels correspond with progressively reduced snow cover within enclosures.

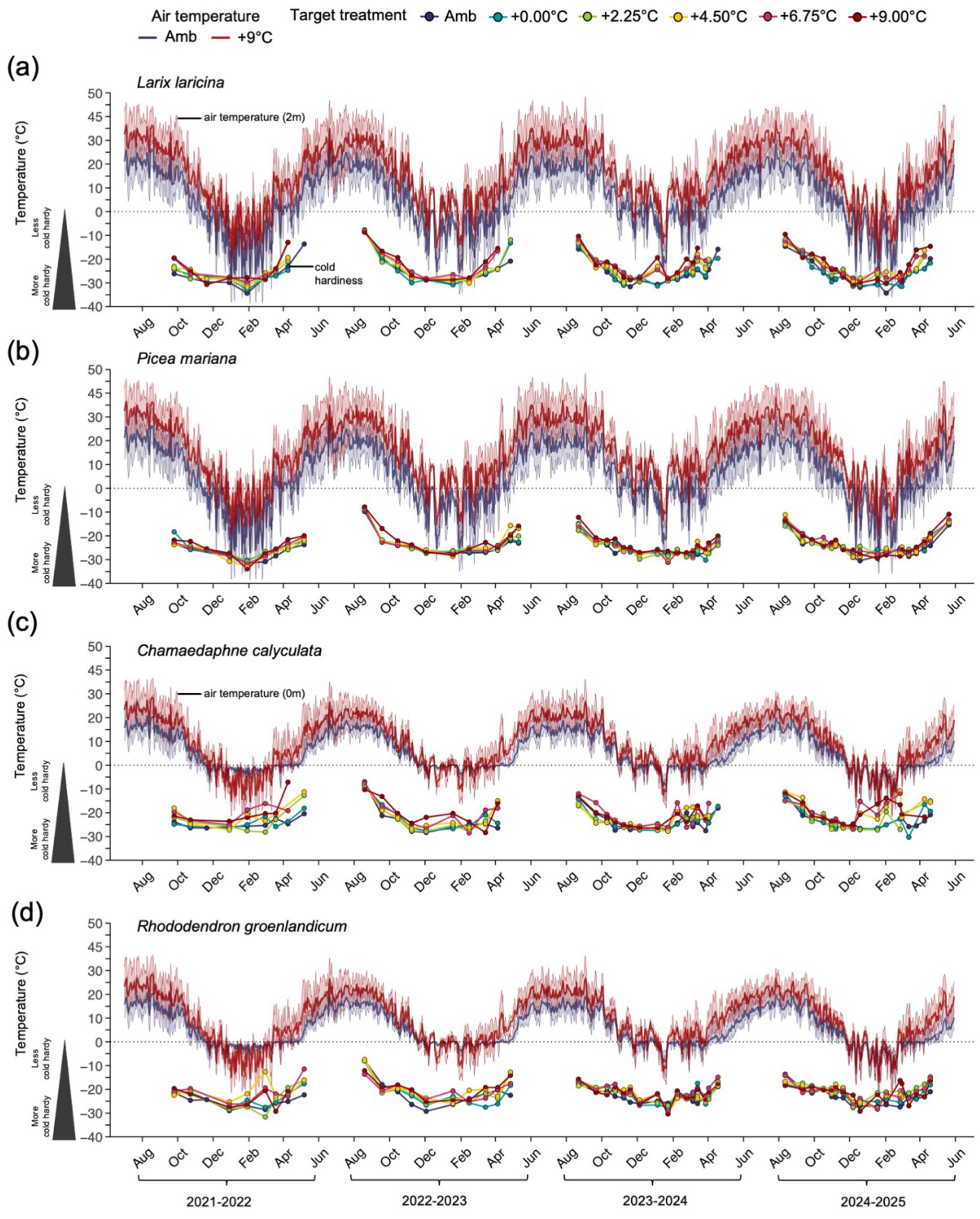


**Figure S3. Seasonal patterns of air temperature and bud cold hardiness across warming treatments.** Seasonal dynamics of air temperature and bud cold hardiness for four species at the SPRUCE experiment from 2021–2025. Panels show (top to bottom) *Larix laricina*, *Picea mariana*, *Chamaedaphne calyculata*, and *Rhododendron groenlandicum*. Lines represent daily air temperature measured within ambient (blue) and warmed (+9 °C; red) treatments, with shaded bands indicating variability across plots. Colored points show measured bud cold hardiness (LTE, °C) across warming treatments (+0.00, +2.25, +4.50, +6.75, and +9.00°C above ambient) at each sampling date during the dormant season.

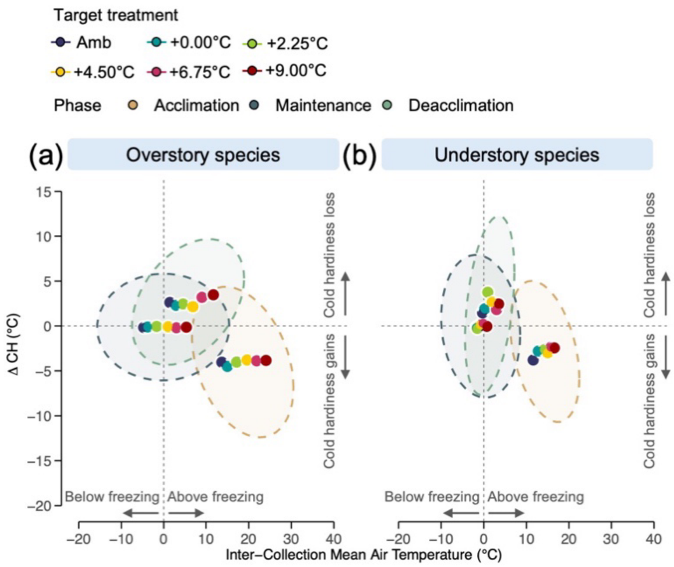


**Figure S4. Short-term thermal responsiveness of cold hardiness across physiological phases and treatments.** Relationships between inter-collection mean air temperature and short-term changes in bud cold hardiness (ΔCH) for overstory (**a**) and understory species (**b**). Negative ΔCH indicates acclimation; positive values indicate deacclimation. Points represent treatment means and ellipses represent 90% confidence regions for each physiological phase: acclimation (golden), midwinter maintenance (dark teal), and deacclimation (sage green).

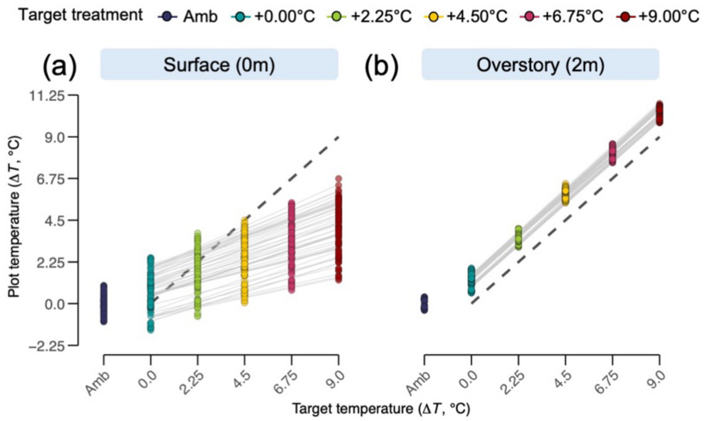


**Figure S5. Cumulative intra-seasonal corrected plot temperature differential (ΔT, °C) versus target warming treatment (ΔT, °C).** Data shown at two sensor heights: (**a**) surface level (0m) and (**b**) overstory (2m). Each point represents the mean cumulative intra-seasonal corrected ΔT for an individual collection and treatment (Ambient, +0.00°C, +2.25°C, +4.50°C, +6.75°C, and +9.00°C). Gray lines connect collection means across treatments within each panel. The dashed line indicates the 1:1 relationship between target and corrected temperatures.

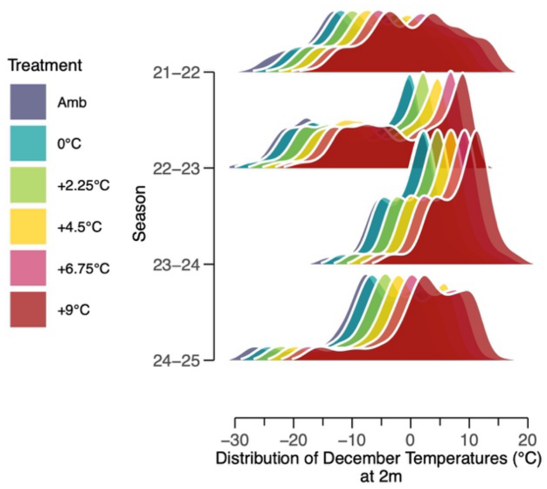


**Figure S6. Histogram of measured December temperatures** for each of four seasons at 2 m at the SPRUCE experiment.

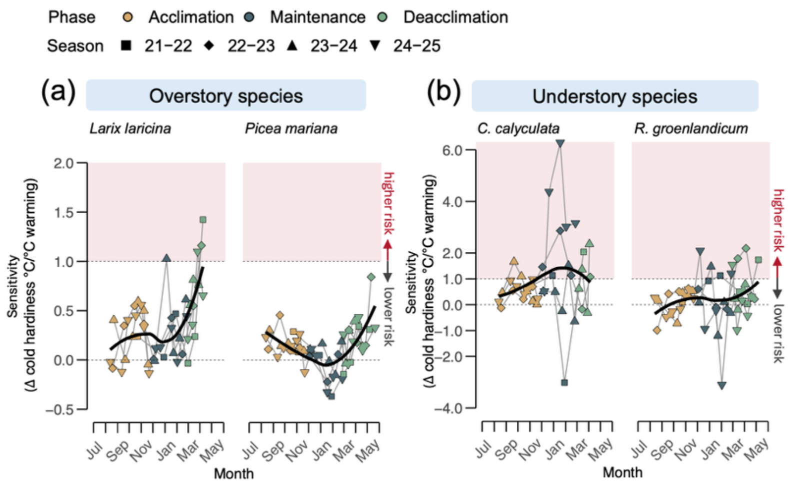


**Figure S7. Warming sensitivity of bud cold hardiness across seasonal phases.** Slopes of warming responses for (**a**) overstory species and understory (**b**) species across acclimation (golden), maintenance (dark teal), and deacclimation (sage green) phases. Each point represents the slope of cold hardiness change per °C warming using inter-collection mean temperatures for an individual collection across four winters. Black curves show the overall seasonal trend; thin grey lines connect points within individual seasons.

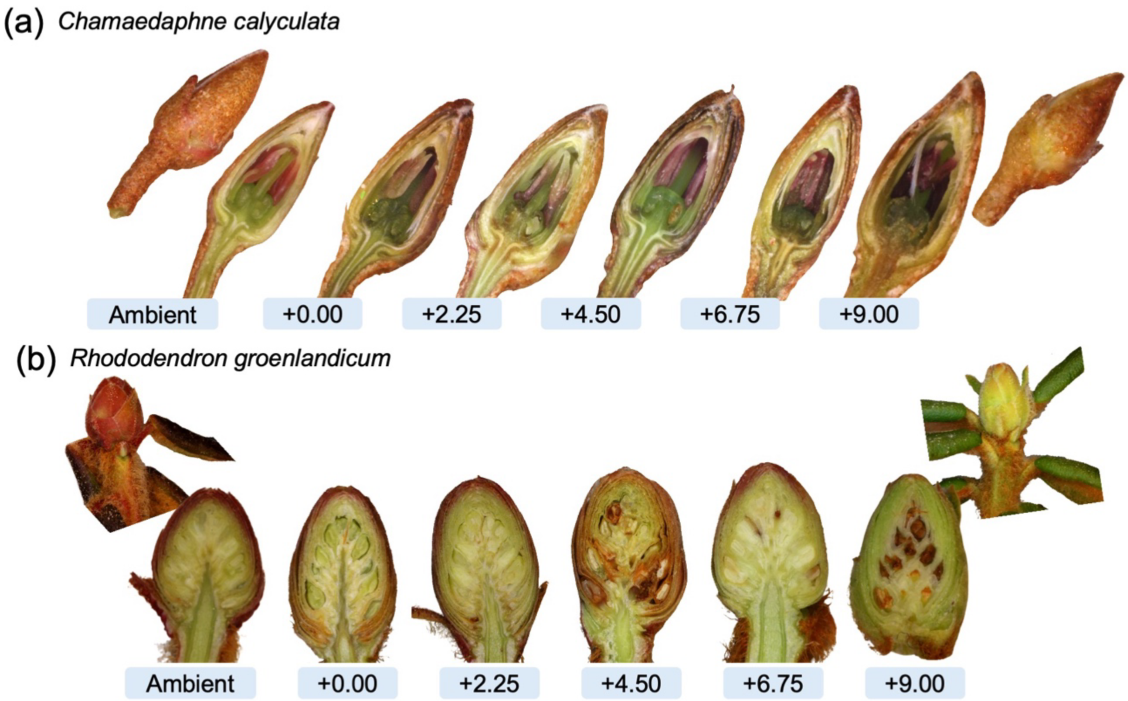


**Figure S8. Bud cross-sections of understory shrub species across warming treatments.** Representative bud cross-sections of (**a**) *Chamaedaphne calyculata* (leatherleaf) and (**b**) *Rhododendron groenlandicum* (Labrador tea) across ecosystem warming treatments. Buds are shown from ambient conditions and warming levels of +0.00°C, +2.25°C, +4.50°C, +6.75°C, and +9.00°C above ambient. Images illustrate structural differences in bud tissues collected in early spring (25 March 2025).

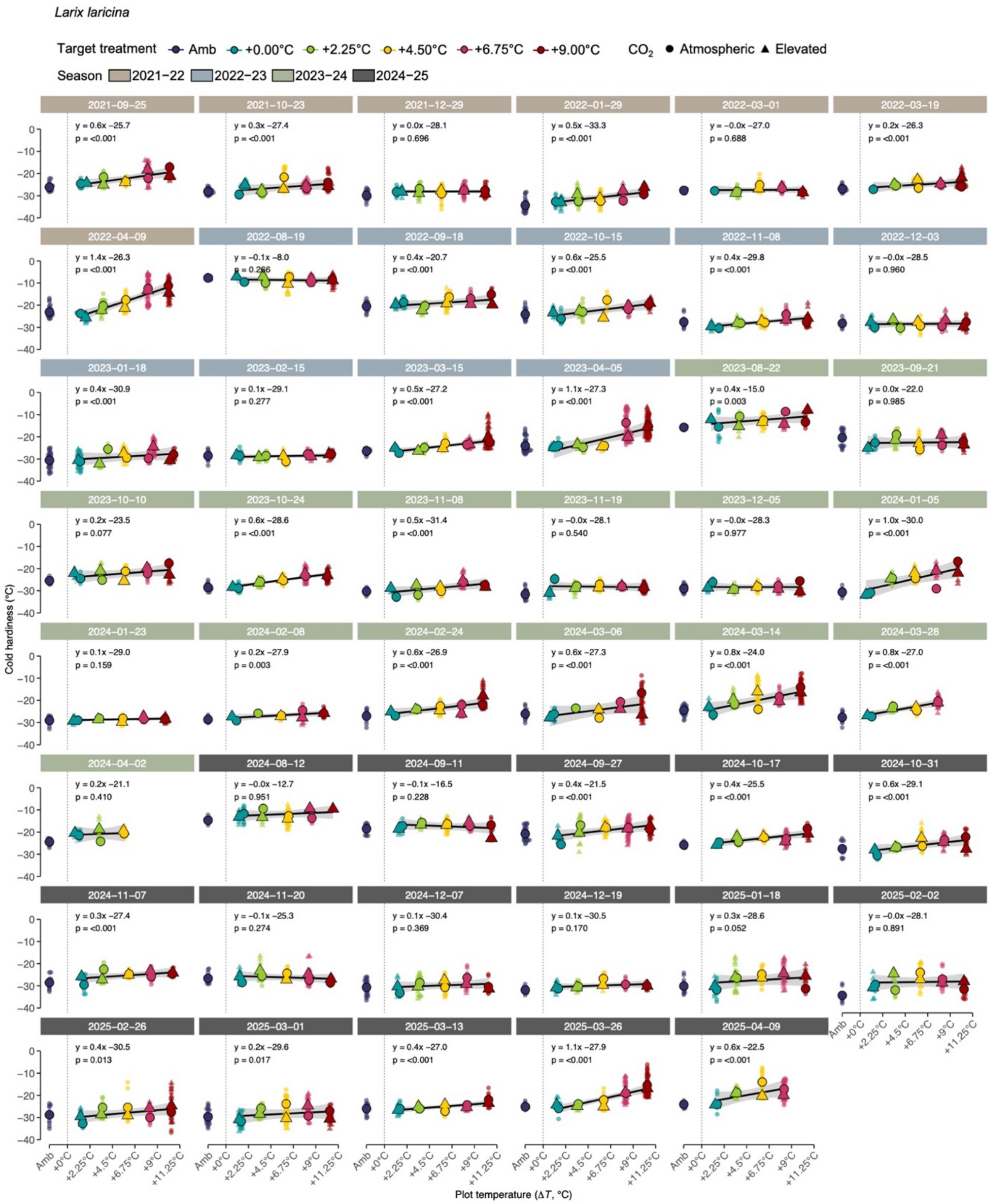


**Figure S9. Seasonal dynamics of bud cold hardiness in *Larix laricina* across warming treatments.** Bud cold hardiness (LTE, °C) measured at multiple sampling dates from 2021–2025 at the SPRUCE experiment. Each panel represents a sampling date, with points showing individual plot measurements across ecosystem warming treatments (+0.00, +2.25, +4.50, +6.75, and +9.00°C above ambient). Colors indicate warming levels. Solid black lines show linear regression relationships between bud cold hardiness and air temperature across treatments for each sampling date.

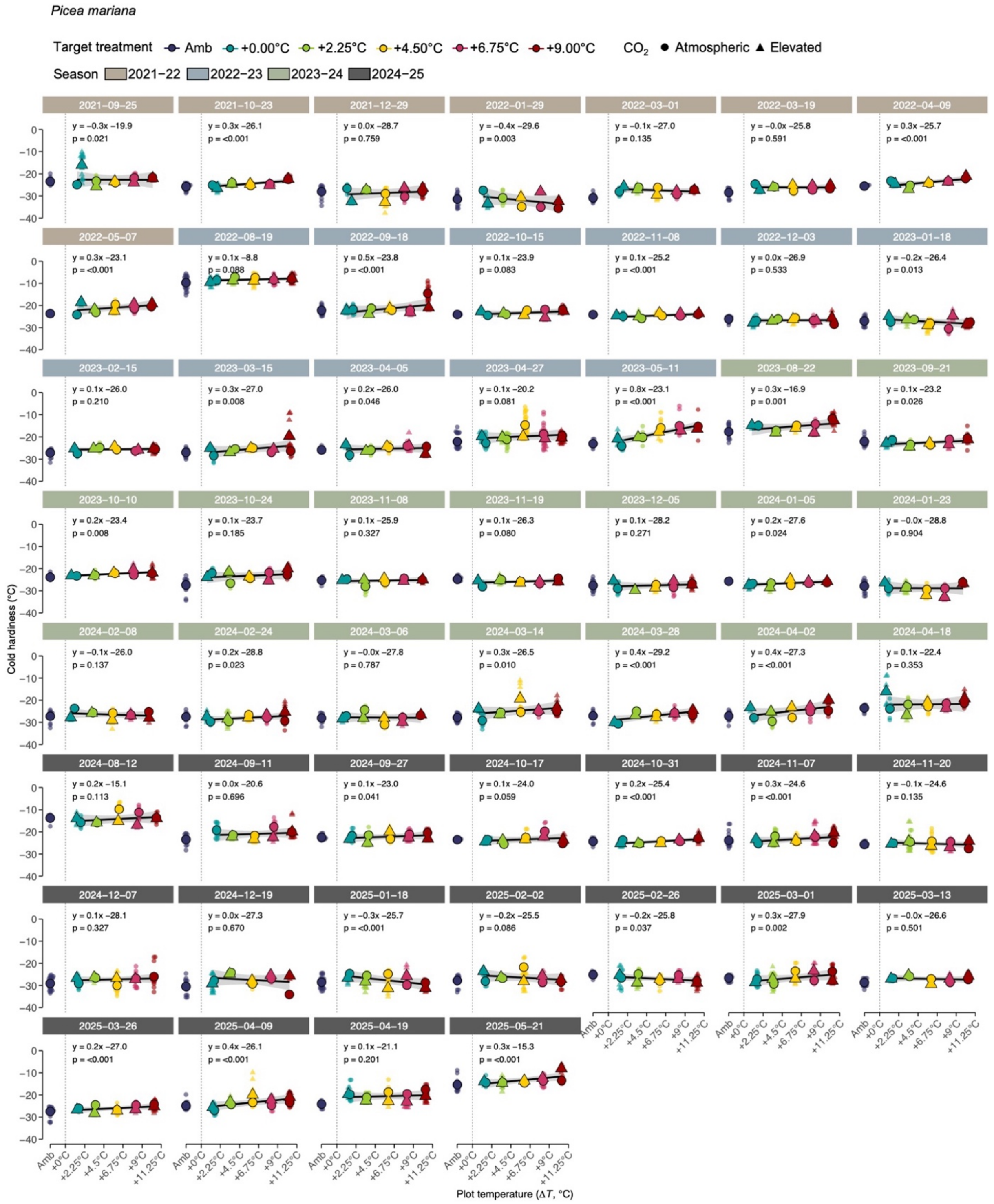


**Figure S10. Seasonal dynamics of bud cold hardiness in *Picea mariana* across warming treatments.** Bud cold hardiness (LTE, °C) measured at multiple sampling dates from 2021–2025 at the SPRUCE experiment. Each panel represents a sampling date, with points showing individual plot measurements across ecosystem warming treatments (+0.00, +2.25, +4.50, +6.75, and +9.00°C above ambient). Colors indicate warming levels. Solid black lines show linear regression relationships between bud cold hardiness and air temperature across treatments for each sampling date.

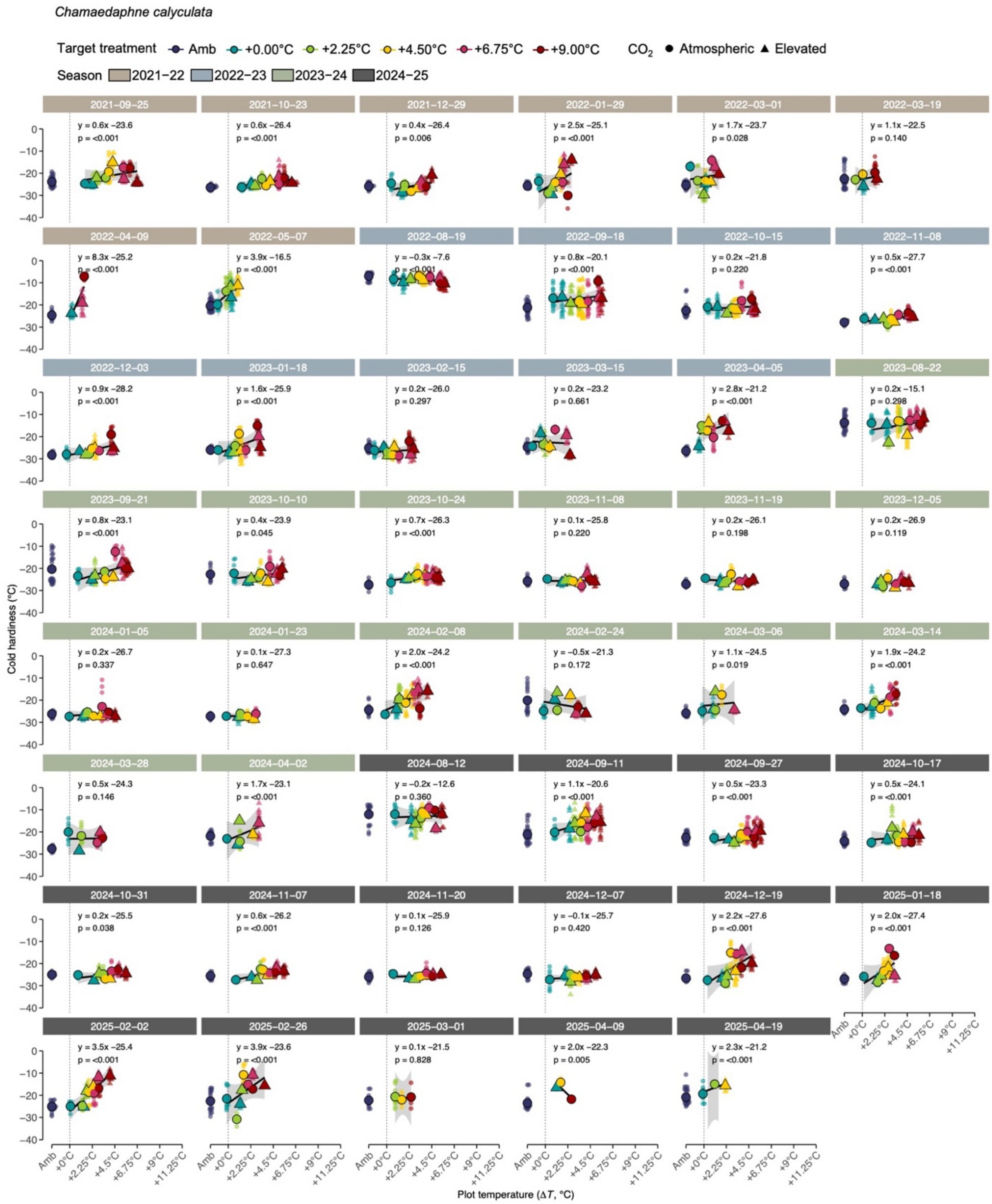


**Figure S11. Seasonal dynamics of bud cold hardiness in *Chamaedaphne calyculata* across warming treatments.** Bud cold hardiness (LTE, °C) measured at multiple sampling dates from 2021–2025 at the SPRUCE experiment. Each panel represents a sampling date, with points showing individual plot measurements across ecosystem warming treatments (+0.00, +2.25, +4.50, +6.75, and +9.00°C above ambient). Colors indicate warming levels. Solid black lines show linear regression relationships between bud cold hardiness and air temperature across treatments for each sampling date.

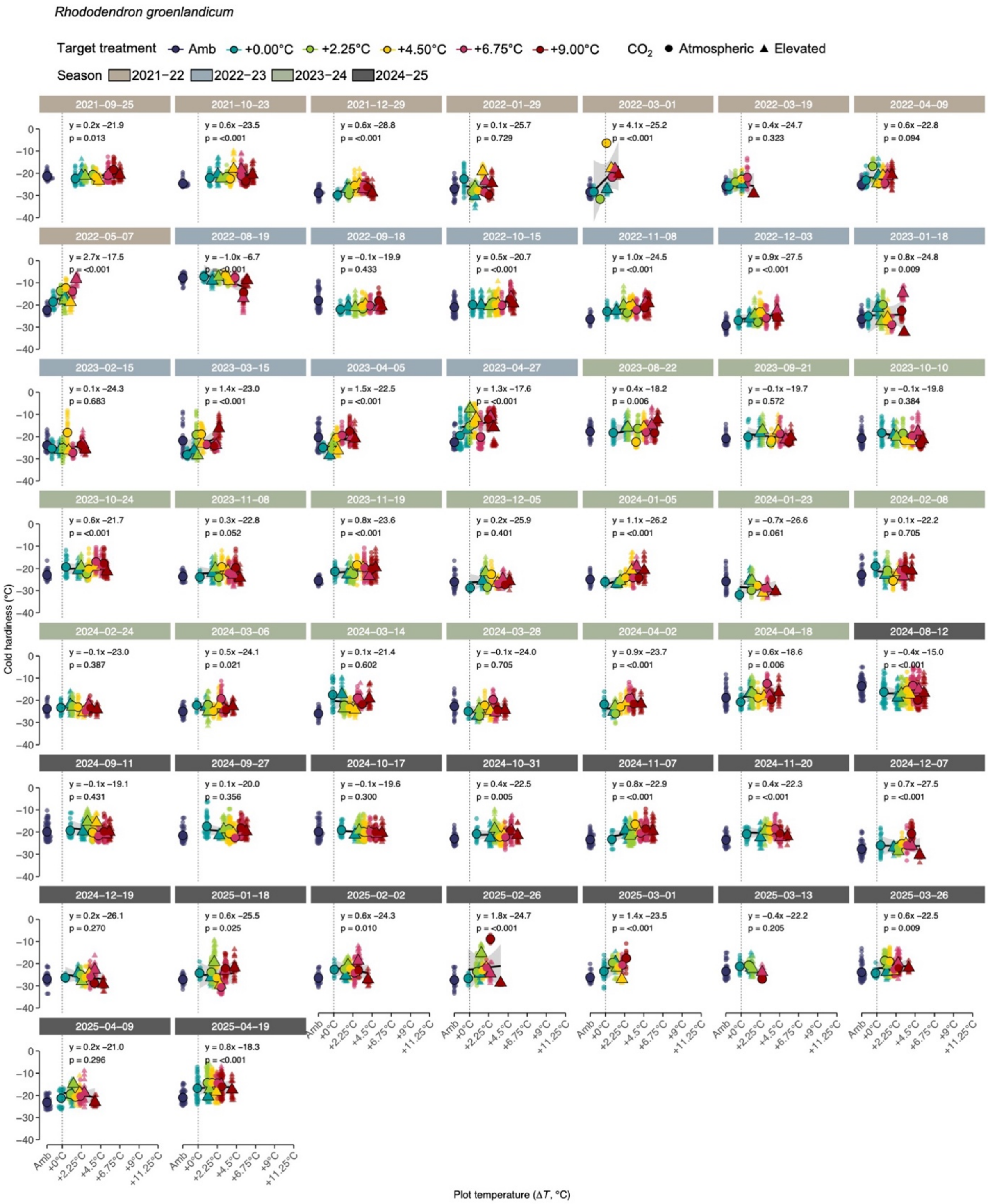


**Figure S12. Seasonal dynamics of bud cold hardiness in *Rhododendron groenlandicum* across warming treatments.** Bud cold hardiness (LTE, °C) measured at multiple sampling dates from 2021–2025 at the SPRUCE experiment. Each panel represents a sampling date, with points showing individual plot measurements across ecosystem warming treatments (+0.00, +2.25, +4.50, +6.75, and +9.00°C above ambient). Colors indicate warming levels. Solid black lines show linear regression relationships between bud cold hardiness and air temperature across treatments for each sampling date.

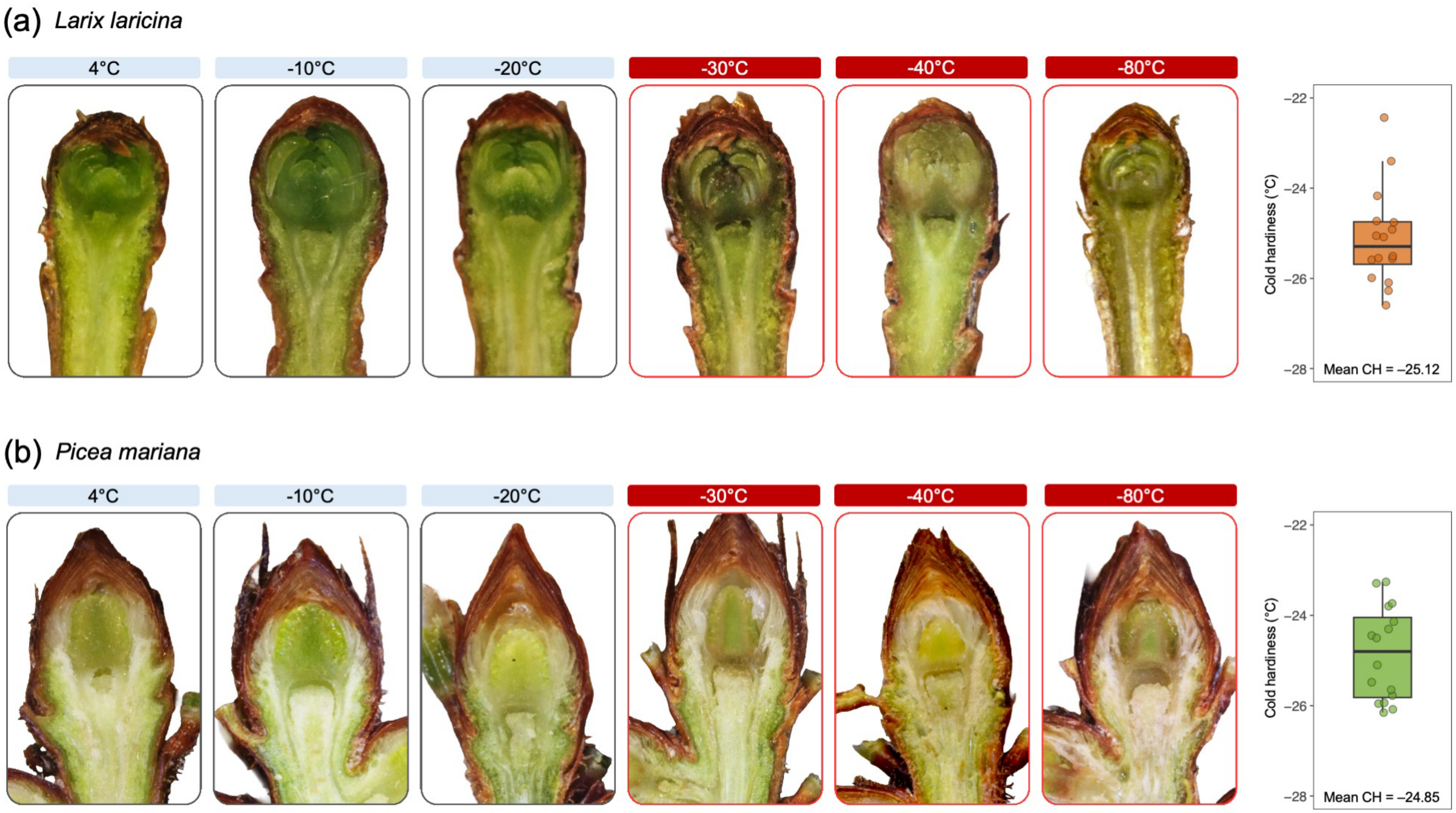


**Figure S13. Control freeze test of overstory species under ambient conditions.** Representative bud cross-sections of (**a**) *Larix laricina* and (**b**) *Picea mariana* exposed to a series of freeze test temperatures (4°C to -80°C). Images illustrate structural differences and visual browning damage in bud tissues tested in early spring (2 April 2025 for *Larix laricina* and 16 April 2026 for *Picea mariana*). Red borders and labels indicate temperatures at which visible tissue damage is observed. Box plots show the distribution of cold hardiness values and individual points represent single bud cold hardiness measurements. Mean cold hardiness was −25.1°C for *Larix laricina* and −24.9°C for *Picea mariana*, consistent with the bud browning observed in the dissections for both species.

**Supplementary Table**

**Table S1. Summary of linear effects model results for cold hardiness across four species.** Linear models evaluating the effects of warming treatment (ΔT), $CO_2$, and collection period on cold hardiness. Values shown are *p*-values from sequential ANOVA tests. Adjusted $R^2$ represents the proportion of variance explained by fixed effects.

| Species | Model | ΔT | $CO_2$ | ΔT × $CO_2$ | Collection | ΔT × Collection | $CO_2$ × Collection | ΔT × $CO_2$ × Collection | Adj. $R^2$ | AIC |
|---|---|---|---|---|---|---|---|---|---|---|
| *L. laricina* | m_lm1 | <0.001 | 0.70 | 0.06 | <0.001 | <0.001 | 0.51 | 0.92 | 0.86 | 2141.79 |
| | m_lm2 | <0.001 | 0.69 | – | <0.001 | <0.001 | – | – | 0.86 | 2077.02 |
| | m_lm3 | <0.001 | – | – | <0.001 | <0.001 | – | – | 0.86 | 2075.34 |
| *P. mariana* | m_lm1 | <0.001 | 0.55 | 0.83 | <0.001 | 0.02 | 0.02 | 0.11 | 0.86 | 2231.39 |
| | m_lm2 | <0.001 | 0.57 | – | <0.001 | 0.06 | – | – | 0.86 | 2220.21 |
| | m_lm3 | <0.001 | – | – | <0.001 | 0.06 | – | – | 0.86 | 2218.56 |
| *C. calyculata* | m_lm1 | <0.001 | 0.25 | 0.80 | <0.001 | <0.001 | 0.03 | 0.04 | 0.82 | 2120.46 |
| | m_lm2 | <0.001 | 0.28 | – | <0.001 | <0.01 | – | – | 0.82 | 2132.96 |
| | m_lm3 | <0.001 | – | – | <0.001 | <0.01 | – | – | 0.82 | 2132.88 |
| *R. groenlandicum* | m_lm1 | <0.01 | 0.81 | 0.40 | <0.001 | 0.15 | 0.79 | 0.20 | 0.77 | 2496.41 |
| | m_lm2 | <0.01 | 0.81 | – | <0.001 | 0.15 | – | – | 0.77 | 2442.61 |
| | m_lm3 | <0.01 | – | – | <0.001 | 0.15 | – | – | 0.77 | 2440.67 |
